# Glioma-associated fibroblasts promote glioblastoma resistance to temozolomide through CCL2-CCR2 paracrine signaling

**DOI:** 10.1101/2024.03.05.581575

**Authors:** Mingrong Zuo, Shuxin Zhang, Siliang Chen, Yufan Xiang, Yunbo Yuan, Tengfei Li, Wanchun Yang, Zhihao Wang, Yuze He, Wenhao Li, Wentao Feng, Ni Chen, Yuan Yang, Yunhui Zeng, Qing Mao, Mina Chen, Yanhui Liu

## Abstract

Complicated tumor microenvironment contributes mostly to chemoresistance in glioblastoma. Glioma-associated fibroblasts (GAFs) were recently identified as non-tumor stromal cells in the glioblastoma microenvironment, whereas their function in glioblastoma chemoresistance is unclear. Herein, we interrogated the correlation between GAFs and chemoresistance of glioblastoma by examining a series of patient-derived GAFs and glioblastoma organoids (GBOs), revealing that GAFs could promote temozolomide resistance in glioblastoma. Mechanistically, C-C motif chemokine ligand 2 (CCL2) secreted by GAFs selectively activated the ERK1/2 signaling in glioblastoma cells to potentiate temozolomide resistance. Pharmacologically disrupting the CCL2-CCR2 axis or MEK1/2-ERK1/2 pathway effectively improved the therapeutic efficacy of temozolomide in GBM cells and patient-derived GBOs, and both decreased phospho-ERK1/2 expression. Collectively, our results identified that targeting the GAF-dominated CCL2-ERK1/2 pathway may be an alternative strategy to alleviate the GAF-mediated chemoresistance of glioblastoma.

**Significance:** Comprehensive interpretation of the mutual support between tumor microenvironment and cancer cells is demanded for glioma with poor response rates to chemotherapy. This study demonstrates that GAFs promote the temozolomide resistance of glioblastoma by secreting cytokine CCL2 to activate ERK1/2 pathway, which may serve as a potential druggable candidate.

Graphic abstract.
Schematic illustration for GAFs mediated chemoresistance of GBMs and underlying mechanisms. We demonstrate that Glioma-associated Fibroblasts (GAFs) grow in gliomas by isolating and identifying a panel of patient-derived GAFs. CCL2 secreted by GAFs stimulates CCR2 in GBM cells, which promotes activation of the ERK1/2 expression to potentiate GBM chemoresistance.

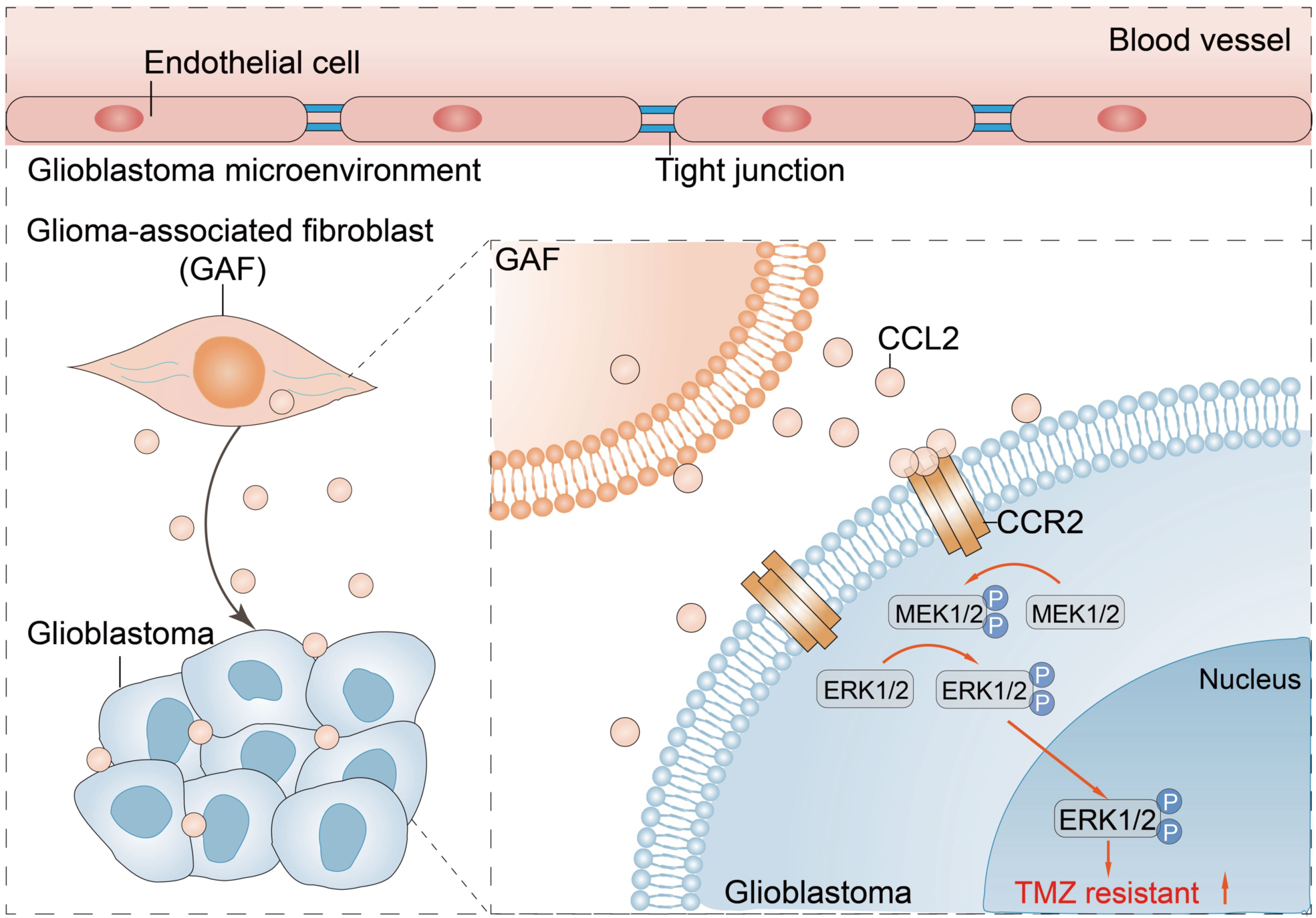

## Introduction

Glioma is the most prevalent primary tumor in brain parenchyma, characterized by its malignant aggressiveness and pronounced lethality ^1,2^. Despite advances in multimodality therapy incorporating maximal safe resection, radiotherapy, and the alkylating agent temozolomide (TMZ) chemotherapy, the clinical outcome of glioma patients remains unsatisfactory, especially for the deadliest type of glioblastoma (GBM) ^3^. Accumulating evidence has associated the poor prognosis of gliomas with their multifaceted microenvironment, which could promote resistance to TMZ and accelerate tumor progression ^4^.

In the tumor microenvironment (TME), non-tumor cells can interact with each other as well as the tumor cells to create a protective niche ^5^. Among all non-tumor cells that settle in the TME, cancer-associated fibroblasts (CAFs) grow in multiple lethal tumors and regulate a variety of malignant characteristics of tumors, such as proliferation, treatment resistance, invasion, and metastasis ^6–12^. The single-cell RNA-sequencing (scRNA-seq) and novel genetically engineered mouse models not only deepen our insight into the phenotypical and functional heterogeneity of CAFs but also shed light on CAF-targeting therapies for tumor treatment ^5^.

A long-standing presumption is that brain tumors may lack CAFs due to scarce fibroblasts in the central nervous system (CNS). However, a fresh study suggests that resident fibroblasts in the CNS are the primary contributors to forming a fibrotic scar in response to inflammation, leading to accentuated illness, which implies that CAFs may exist in brain tumors ^13^. A pilot bioinformatic study analyzed the Chinese Glioma Genome Atlas, demonstrating that the higher proportion of CAFs independently correlates with poor outcomes among glioma patients ^14^. In addition, scRNA-seq analyses recently revealed the presence of stromal cells expressing CAF markers in GBMs, identified as CAFs with pro-tumoral effects through interacting with GBM stem cells and inducing M2 macrophage polarization ^15,16^. Despite the critical role of CAFs in the progression of numerous solid tumors, the lack of appropriate methodology for isolating pure patient-derived CAFs from glioma tissues restrains further exploration of the interaction between CAFs and glioma cells ^15^. Despite the above scRNA-seq findings that GAFs grow in GBMs, the concrete in vitro and in vivo experimental evidence elucidating the existence of GAFs and their biological functions in glioblastoma chemoresistance remains insufficient.

Given these backgrounds, the present study achieved the isolation and expansion of primary glioma-associated fibroblasts (GAFs) from glioblastomas and low-grade gliomas through our optimized approach based on the conventional method ^17^. We confirmed their identity as cancer-associated fibroblasts, termed GAFs, by comprehensively analyzing their molecular characteristics using RNA transcriptomes and scRNA-seq. We further interrogated their role in the context of GBM resistance to TMZ by in vitro and ex vivo GBM organoid (GBO) models. Our results suggested that GAFs secreting cytokine CCL2 potentiate TMZ resistance of GBM cells through CCL2-CCR2 mediated ERK1/2 signaling activation. These findings revealed the protective role of GAFs in GBM chemoresistance, thereby serving as a candidate for medical intervention.

## Results

### GAF informs poor chemotherapeutic response for glioma patients

GAFs have been identified recently as a new kind of stromal cell in the GBM microenvironment ^15^. However, whether GAFs populate in low-grade gliomas (LGGs) remains unelucidated. Therefore, we first analyzed another scRNA-seq dataset (GSE182109) with 18 glioma samples (2 LGGs, 11 primary GBMs, and 5 recurrent GBMs) published by Abdelfattah N et al to further validate the presence of GAFs in distinct grades of gliomas ^18^. After sample integration and dimension reduction, high-quality single cells in the whole dataset were aggregated into 19 clusters; Most cells in cluster 12 were annotated as fibroblasts by SingleR (Figures 1A and S1A). In line with GBM samples in the dataset (Figure S1B), GAFs could also be annotated in the LGGs (Figure S1C). Copy number variation (CNV) profile of clusters showed that these cells had lower sums of scale CNV proportion in their genome than the astrocytic clusters, which suggested they were non-malignant cells (Figures S1D and S1E). Differential expression analysis found that these cells expressed significantly higher levels of known CAF markers (Fibronectin (FN1), TAGLN, ACTA2, and TPM2) ^19^ (Figure S1F). Notably, FN1 and ACTA2 could separate cluster 12 from other clusters (Figures S1G and S1H). Our single-cell results confirmed the presence of GAFs in distinct grades of the glioma microenvironment.

**Figure 1.**
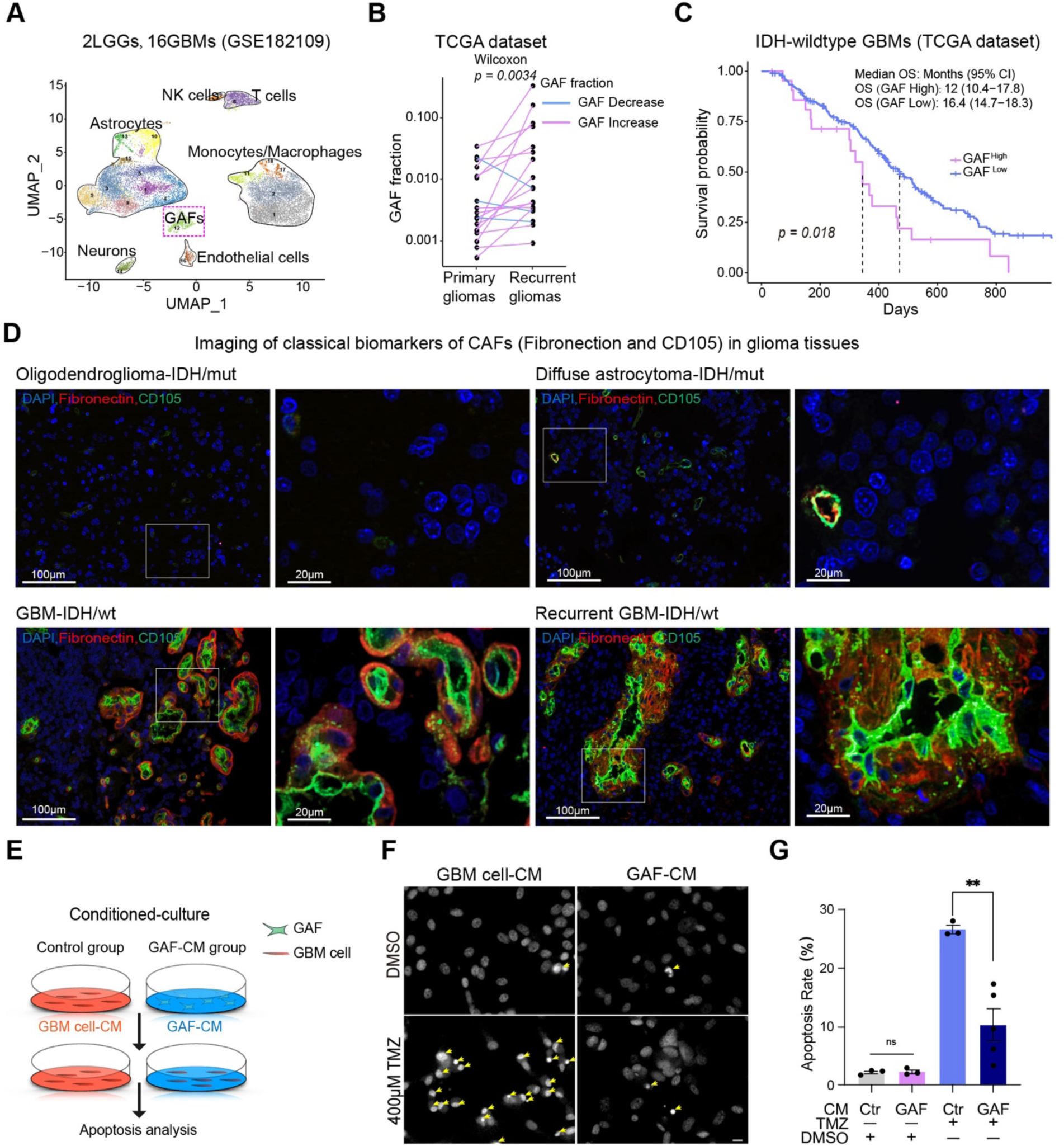
GAFs are associated with worse survival and poor TMZ response for gliomas. (A) UMAP plot presentation of cells clustered by Seurat and their SingleR annotation based on the GSE182109 scRNA-seq dataset (2 LGGs, 11 primary GBMs, and 5 recurrent GBMs). (B) The variation of GAF proportion in the primary and recurrent gliomas from the TCGA database. (C) Kaplan–Meier survival analysis of overall survival of patients with IDH-wildtype GBMs from the TCGA database. (D) Immunofluorescence staining of CD105 (Green) and Fibronectin (Red) in the primary IDH-wildtype GBM (GBM-IDH/wt), IDH/wt recurrent GBM, IDH mutant (IDH/mut) oligodendroglioma, and IDH/mut diffuse astrocytoma. (E) Schematic diagram of conditioned culture. GBM cells were cultured in GAF-CM (Experiment group) and GBM cell-CM (Control group) respectively for indicated hours and then treated by DMSO or TMZ (400µmol/L). (F) Hochest33258 staining of tumor cell apoptosis after indicated treatment. Scar bar, 20µm. (G) Quantification of apoptotic GBM cells. (n > 3 per group). Data in B, C, and G were mean±SEM. p values from the two-tailed Wilcoxon ranked sum test (B), log-rank test (C), and two-tailed student-t-test (G).

We next analyzed the association of GAFs with clinical characteristics of gliomas. Firstly, the computationally estimated GAF fraction by EPIC R package arose with the histological grading of gliomas (Figure S1I). The GAF fraction was significantly higher in isocitrate dehydrogenase /Wildtype (IDH/WT) GBMs than in the IDH/Mutant gliomas from the TCGA cohort (Figure S1J). We further compared the fraction of GAFs in GBMs and LGGs based on the single-cell dataset ^18^, revealing that GAF (cluster 12) fractions in GBMs (IDH/WT, n=20) were significantly higher than LGGs (IDH/Mutant, n=4) (Figure S1K). Additionally, the computationally estimated GAF fraction increased significantly along with tumor recurrence (Figure 1B). Within each IDH subtype, a higher GAF fraction was likely associated with poorer overall survival (Figures 1C and S1L). These findings implied that GAF may be associated with treatment failure, for example, promoting drug resistance. We further performed immunofluorescence of human glioma tissues for CAF markers, such as FN1 and CD105 ^6,20^, revealing that the primary GBMs and recurrent GBMs were vastly enriched for FN1 and CD105, whereas oligodendrogliomas and diffuse astrocytomas expressed scarce signals of these GAF markers (Figure 1D). These data demonstrated that GAFs were potential risk contributors to the lethality of gliomas. As S. Jain had shown that GAFs could promote GBM growth and induce M2 macrophage polarization ^15^, whether GAFs contributed to the therapeutic failure remains unknown. Thus, we tried to examine the response of GBM cells upon TMZ treatment following GAF conditioned medium (GAF-CM) culture for 48 hours (Figure 1E). Hochest33258 staining found a 16.2% decrease in the apoptosis rate of U87 cells cultured with GAF-CM, compared to the control group (Figures 1F and 1G). Taken together, we put forward that GAFs may drive increased TMZ resistance of GBM cells.

### Isolation and identification of patient-derived GAFs from human gliomas

TME has emerged as an indispensable contributor to intertumoral heterogeneity and treatment resistance of glioma ^21^. Although scRNA-seq analysis recently identified that GAFs existed in GBMs ^15^, a rapid and effective primary cell culture method has not yet been developed. We first generated patient-derived GAFs from glioma patients based on the traditional method of isolating primary CAFs ^17^ following our optimization (Figure 2A). The cell culturing procedure included two vital steps, we first cultured digested cell mixtures of glioma tissues using the neural stem cell medium for 24 hours to separate suspended growth cells, such as cancer stem cells and neural stem cells, from adherent cells. Then, the cell medium containing all non-adherent cells was removed and adherent cells were kept to further derive patient-derived cells (PDCs). In total, thirteen different strains of PDCs from seven lower-grade gliomas and six GBM samples were established. The patient demographic information is listed in Table S1.

**Figure 2.**
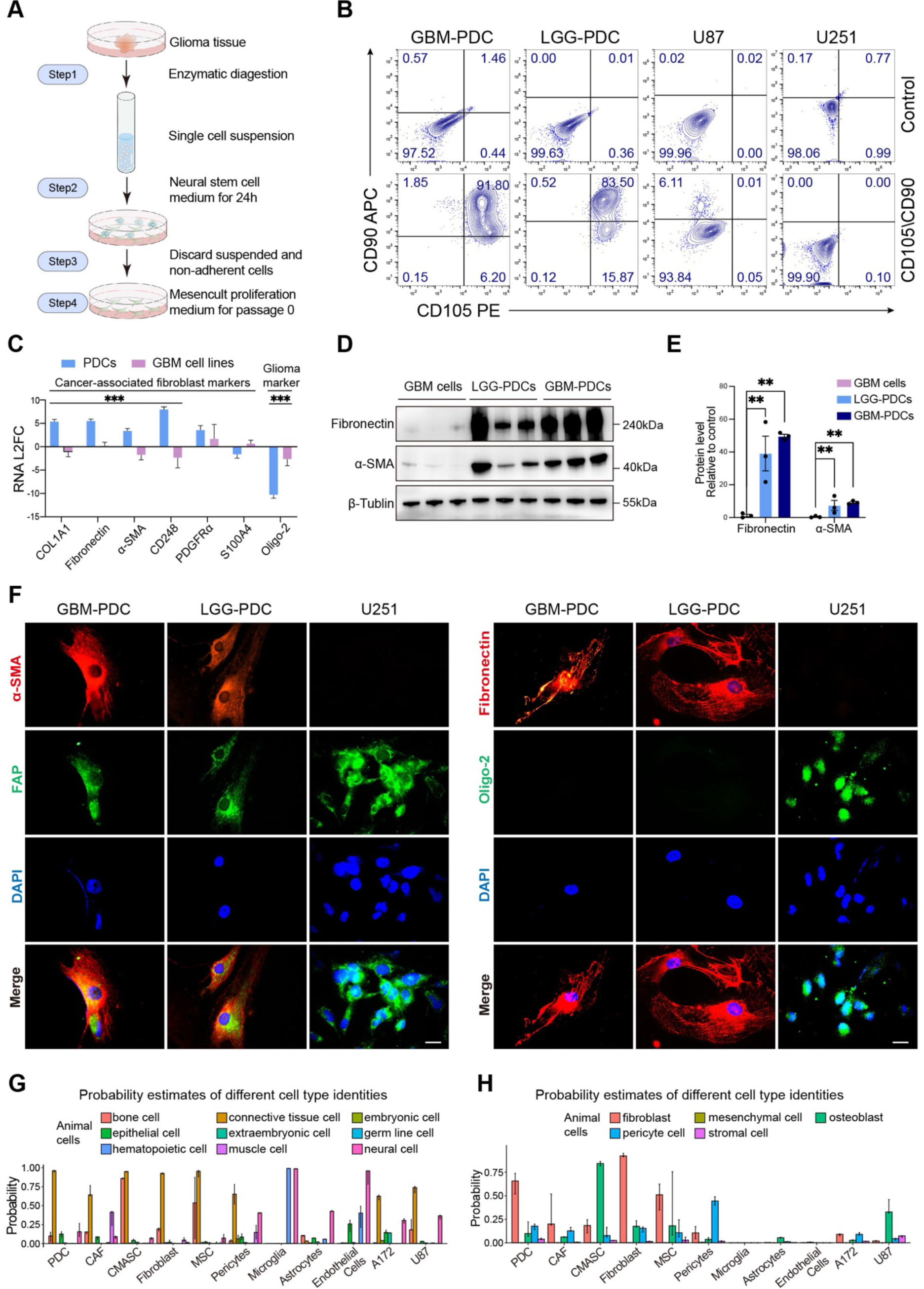
Primary culture and identification of patient-derived GAFs from human gliomas. (A) Schematic diagram of the optimized screening and culturing method of primary glioma-associated fibroblasts from glioma tissues. (B) Proportions of PDCs (PDC01, 08) expressing CAF markers (CD105, CD90) were analyzed by flow cytometry (FCM). Images of the upper row were blank control without primary antibodies staining. (C) mRNA levels of canonical CAF markers (COL1A1, Fibronectin, α-SMA, CD248, PDGFRα, S100A4) and glioma marker (Oligo-2) in PDCs (PDC01, 02, 09, 10, and 11) and glioma cell lines (U87, U251, and A172) quantified by qRT-PCR. (D) Protein levels of CAF markers in PDCs (01, 02, 08, 09, 10, and 13) and glioma cell lines (A172, U87, and U251) (E) Quantification of protein levels. (F) Immunofluorescence staining of α-SMA, Fibronectin, FAP, and Oligo-2 of PDCs (PDC02, 12) and U251 cells. Scale bar, 10µm. (G and H) The probability of cell type identities of the PDCs and other cells in the Cell Ontology family of Animal Cells and Connective Tissue Cells. Data in (C) and (E) were mean ± SEM. p values from two-tailed student-t-test (C) and one-way ANOVA (E). ** *P* < 0.01, *** *P* < 0.001.

We conducted a series of biomolecular analyses to identify the lineage of PDCs. PDCs had larger dimensions and heterogeneous morphologies, such as spindle, cruciform, or stellate-shaped, which resembled the morphology of CAFs (Figure S2A) ^6^. Flow cytometry revealed that more than 90% of PDCs expressed biomarkers of CD105 and CD90, which were reported to be expressed in the majority of CAFs ^20,22,23^; However, glioma cells were negative for these CAF markers (Figures 2B and S2B). Additionally, we found that PDCs indeed expressed CAF-specific markers CD248 (Endosialin, a pan-fibroblast marker), COL1A1, and FN1; The non-specific markers α-SMA (ACTA2), PDGFRα, and S100A4 were also found to be highly expressed in PDCs (Figure 2C) ^24,25^.

On the contrary, the classical biomarker of glioma cells, Oligo-2, was prominently downregulated in all PDCs (Figure 2C), indicating PDCs did not arise from neuro-epithelial cells. Similar results were also observed in the Western Blot analysis (Figures 2D and 2E). Immunofluorescence staining further showed that PDCs expressed both α-SMA and FN1; Conversely, only U251 cells expressed Oligo-2 (Figure 2F). As FAP is non-specific for CAF, either PDCs or U251 cells were positive for FAP (Figure 2F). Taken together, we considered glioma patient-derived PDCs as CAF-like stromal cells.

To ascertain the identity of the CAF-like PDCs in an expanded molecular background, we conducted mRNA-sequencing of these patient-derived CAF-like stromal cells. With the transcriptome data, we predicted the probability of the cells being each cell type in the Cell Ontology vocabulary system using pre-trained cell type classification models. The PDCs showed high a probability of being connective tissue cells in the Animal Cells Family (mean probability = 0.969), and of being fibroblasts in the Connective Tissue Cells family (mean probability = 0.640) (Figure S2C). This identification result placed the PDCs under the Connective Tissue Cells— Fibroblast lineage. Similar identity profiles were also observed with previously reported CAFs, bona fide skin fibroblasts, and mesenchymal stem cells (MSCs) (Figures 2G and H). However, compared to PDCs and fibroblasts, the MSCs also showed a higher probability of being Bone Cell, Osteoblast, and Non-terminally Differentiated Cell (Figure 2H), suggesting that MSCs possessed higher multi-pluripotency, while PDCs were at a later stage of cell differentiation. Copy number analysis based on the RNA-seq data was able to discover known large-scale CNV in the glioma cell lines and glioma tissues (Figure S2D). On the contrary, no significant large-scale CNV was detected in the GAFs from either high-grade or low-grade gliomas (Figure S2D). These findings were in line with those of the normal brain tissue (Figure S2D), indicating that the GAFs were diploid stromal cells rather than malignant cells that underwent epithelial-to-mesenchymal transition. Based on these findings, we concluded that the patient-derived CAF-like stromal cells can be identified as GAFs in the microenvironment of gliomas.

### GAF promotes the TMZ resistance of GBM cells in vitro and in vivo

To determine whether GAFs could confer TMZ resistance to GBM cells, we further detected cell apoptosis through flow cytometry (FCM) analysis. It showed a reduction of apoptotic GBM cells following the GAFs-CM culture (Figures 3A and 3B). To understand whether GAFs could promote TMZ-resistance of GBM cells more efficiently through cell coculture, we further cocultured GAFs and GBM cells (Figure S3A). The anti-apoptotic effects of GBM cells upon TMZ treatment were enhanced by 13.3% with GAFs coculture for 48 hours, compared to GBM cells coculture (Figures 3C and 3D). Moreover, we found that the proportion of apoptotic GBM cells plummeted by more than 20% following 96 hours of coculturing with GAFs (Figures S3B-S3E). These findings suggested that the chemoresistance of GBM cells was correlated with the duration of interaction with GAFs. In the U87 cells cocultured with GAFs, we observed a significant reduction in cleaved-caspase3, 9, and BAX in response to TMZ treatment (Figures 3E and S3F), which further confirmed the protective effect of GAFs on GBM against TMZ. Taken together, we demonstrated that GAFs protected GBM cells from TMZ-induced apoptosis in vitro.

**Figure 3.**
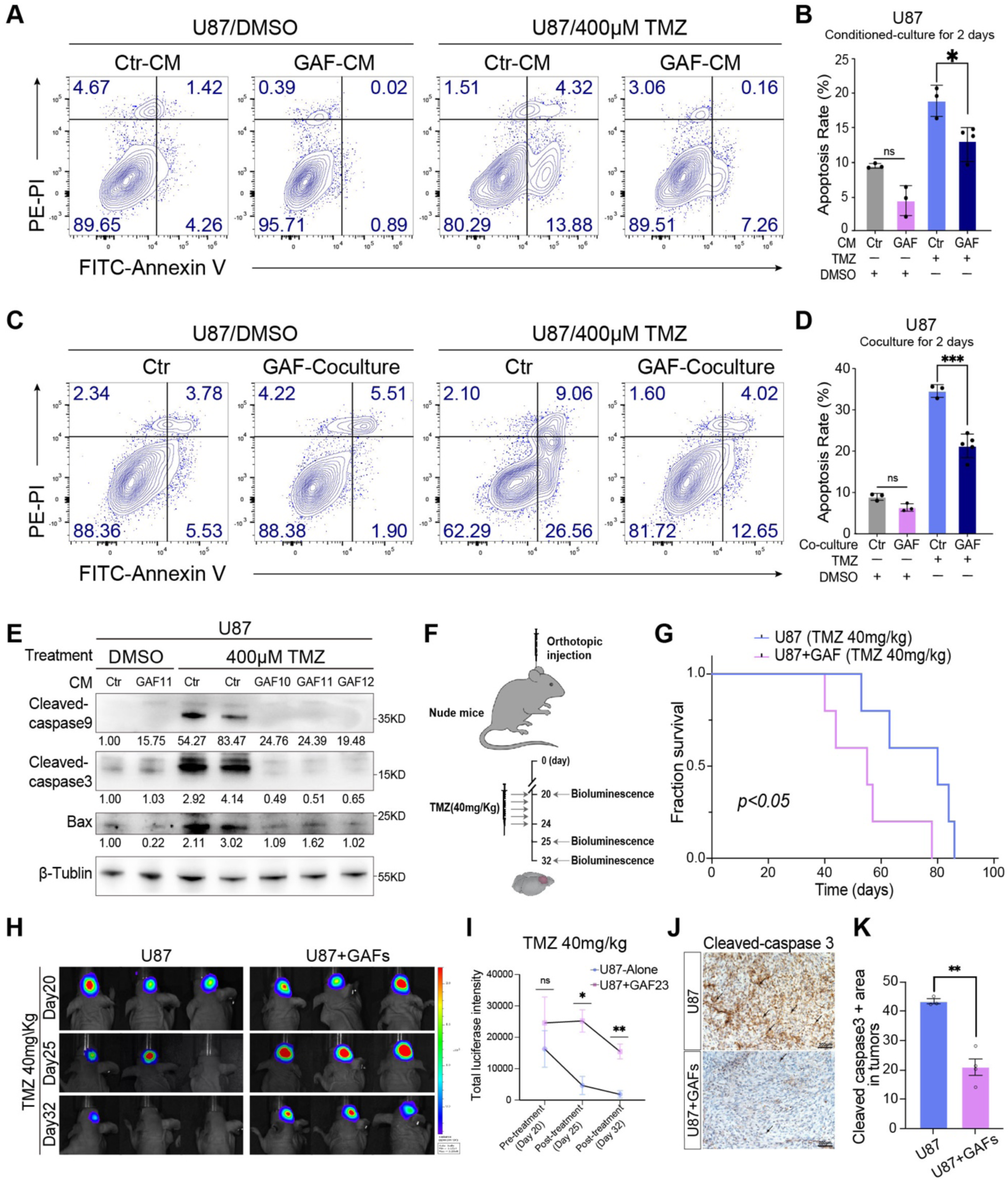
GAF promotes the TMZ resistance of GBM cells in vitro and in vivo. (A) Representative images of the percentage of apoptosis of U87 cells with conditioned culture following TMZ treatment. (B) Quantification of apoptotic GBM cells (n > 3 per group). (C) Representative images of the percentage of apoptosis of U87 cells following 48 hours of cell. coculture. (D) Quantification of apoptotic GBM cells (n > 3 per group). (E) WB analysis of cleaved-caspase9, cleaved-caspase3, and Bax in U87 cells cocultured with. GAFs or U87 cells following indicated treatment. (F) Schematic diagram of orthotopic co-injection of GAFs (1×10^4^) and luciferase-expressing U87 cells (5×10^4^). Twenty days after the orthotopic GBM models were established, mice were treated with TMZ (40mg/Kg, ip, for 5 consecutive days), and tumor growth was monitored through bioluminescence detection at days 20, 25, and 32. (G) Kaplan-Meier survival analysis of mice bearing with U87 alone or U87-GAFs mix orthotopic xenograft treated with TMZ (40mg/Kg) (n=5 per group). (H) In vivo bioluminescence images of xenograft tumor growth during TMZ treatment on days 20, 25, and 32 after the tumor was implanted. (I) Quantification of bioluminescence intensity of tumors. (J) Representative IHC image of cleaved-caspase3 from U87 alone (n = 3 mice per group) and U87+GAFs (n=3 mice per group) following TMZ treatment (K) Quantification cleaved-caspase3 positive areas of tumors. Experiments of A-K were independently used at least three different GAFs. Data in B, D, G, I, and K were mean±SEM. p values from two-tailed student-t-test (B, D, I, and K) and log-rank test (G). * *P* < 0.05, ** *P* < 0.005, *** *P* < 0.001, ns = no significant.

To find out whether GAFs could abate TMZ therapeutic efficacy in vivo, we then challenged the nude mice with an orthotopic co-injection of GAFs and luciferase-expressing U87 cells (Figure 3F). Kaplan-Meier survival analysis revealed that animals with orthotopic GBM-GAFs mix had significantly shorter survival time than GBM alone (Figure 3G). In line with survival analysis, the orthotopic GBM mixed with GAFs displayed minimal response to TMZ. Conversely, the growth of orthotopic GBM alone was efficiently inhibited after treatment (days 25 to 32) (Figures 3H and 3I). IHC staining also showed that the expression of cleaved-caspase 3 reduced significantly in the orthotopic GBMs mixed with GAFs relative to the GBMs alone under TMZ treatment (Figures 3J and 3K). Collectively, our experiments demonstrated the function of GAFs in promoting GBM resistance to TMZ chemotherapy.

### GAFs promote TMZ resistance of GBM cells by preferentially secreting CCL2

To identify what factors participated in the GAF-GBM interaction, we first evaluated the distinguishing expression of the cytokine-encoding genes in GAFs and GBM cells. Among cytokine-encoding genes cataloged in the Cytokine Registry (https://www.immport.org/resources/cytokineRegistry), CCL2 expressed at a significantly higher level in the GAFs compared to GBM cells (Figure 4A). Recently, accumulating evidence suggested that CCL2 not only mediated chemotaxis of several immune cells but also could induce multiple solid tumors resistant to a broad spectrum of anticancer drugs ^26–28^. Correlation analysis showed that GAF was positively associated with the expression level of CCL2 in both IDH-wildtype and IDH-mutant gliomas (Figures 4B and S4A). WB analysis also revealed that GAFs expressed significantly higher CCL2 as compared to GBM cells (Figure 4C). Moreover, ELISA analysis found that all GAFs secreted significantly higher levels of CCL2 compared to the GBM cells (Figure 4D). Given the tumor-protecting role of GAFs and their highly ranked CCL2 expression, we managed to investigate whether CCL2 was the main mediator that linked the GAFs with GBM cells. Firstly, we found human CCL2 indeed exerted a tumor-protecting effect on GBM cells upon TMZ treatment (Figures 4E and 4F). Then, we performed a rescue assay to study whether the CCL2-CCR2 inhibitor would attenuate the GAF-mediated tumor-protecting effect. INCB3344 was found to be a potent, selective, and orally available antagonist of CCR2 ^29^, which could effectively inhibit CCL2-CCR2 signaling. Firstly, WB analysis suggested an incremental preventing effect of INCB3344 against GAF-mediated TMZ resistance (Figure S4B). Treatment with INCB3344 during cell-conditioned culture dramatically inhibited the protective effect of GAFs on TMZ-induced GBM cell apoptosis (Figures 4G and 4H). Consistently, WB analysis confirmed that disrupting the CCL2-CCR2 axis could significantly up-regulate the expression of cleaved-caspase 3 and 9 of GBM cells, suggesting INCB3344 could effectively block the protection from TMZ-induced apoptosis by GAFs (Figures 4I-K). Taken together, our results revealed a paracrine mechanism through which GAFs reduced the chemosensitivity of GBM cells to TMZ by the CCL2-CCR2 axis (Figure 4L).

**Figure 4.**
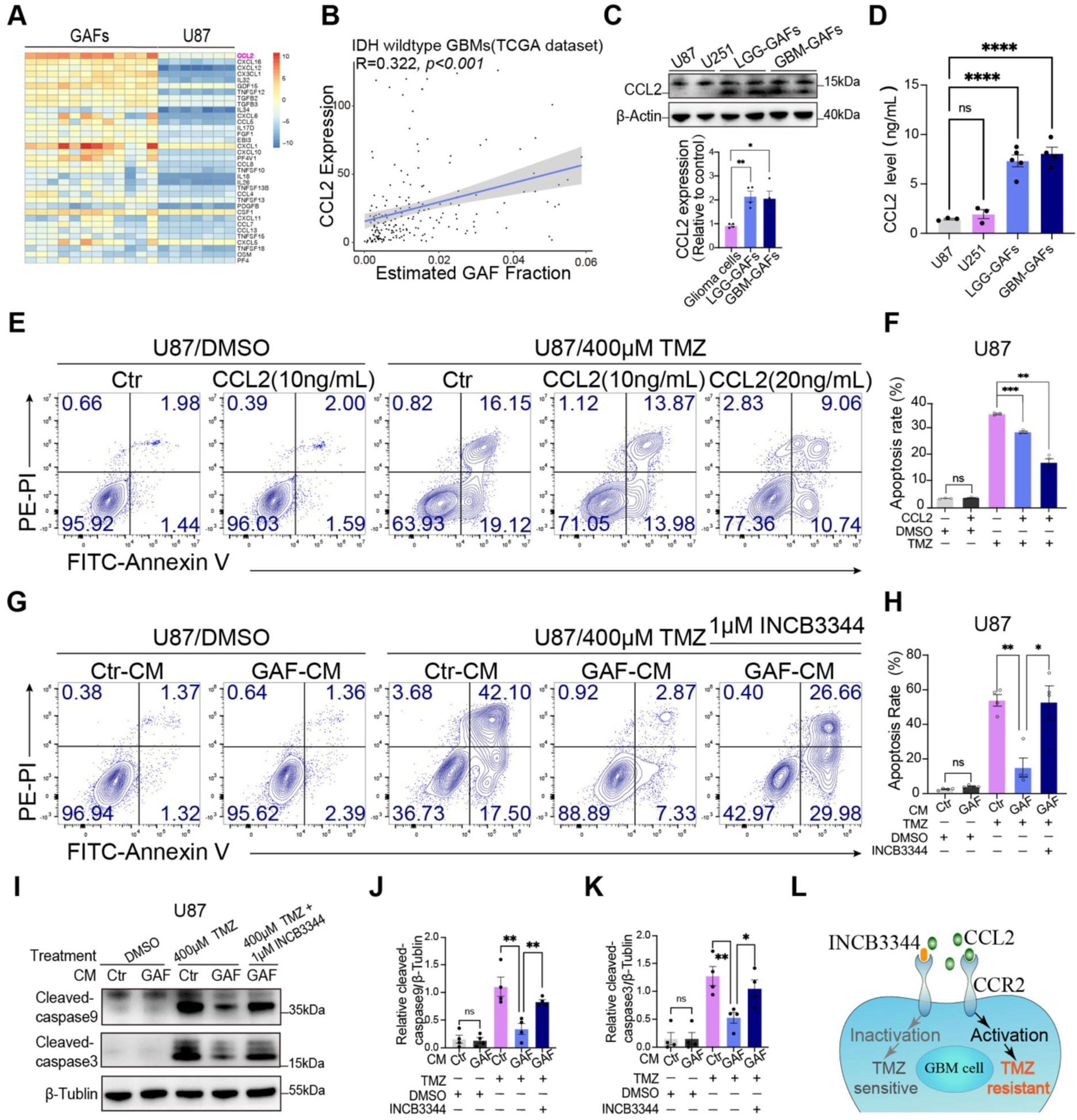
GAF promotes TMZ resistance of GBM cells via cytokine CCL2. (A) Expression of cytokines and chemokines in human GAFs (n = 12 different GAFs) relative to U87 cells (n = 6) identified by RNA-seq. (B) Pearson correlation analysis of CCL2 expression and estimated GAF fraction of GBMs from the TCGA database (n = 218). (C) WB analysis of CCL2 level in GBM cells (U87 and U251), LGG-GAFs, and GBM-GAFs (n = 4 different GAFs for each group). Quantification of CCL2 expression is at the bottom. (D) GBM cells (U87, U251), LGG-GAFs, and GBM-GAFs were cultured with 1% FBS medium for 72 hours. Elisa analysis of CCL2 level in U87-CM (n = 3), U251-CM (n = 3), LGG-GAF-CM (n = 4), and GBM-GAF-CM (n = 3). (E) Representative images of the percentage of apoptosis of U87 cells treated with human CCL2 following TMZ treatment. (F) Quantification of apoptotic GBM cells (n = 3 per group). (G) Representative flow cytometry plots of U87 cells cultured with Ctr-CM, GAF-CM, and GAF-CM mixed with INCB3344 (1µmol/L) for 48 hours following TMZ treatment (H) Quantification of apoptotic GBM cells (n = 4 per group). (I) WB analysis of cleaved-caspase9 and cleaved-caspase3 in U87 cells cultured with Ctr-CM or GAF-CM following indicated treatment. (J and K) Quantifications of protein levels of cleaved-caspase9 and cleaved-caspase3 respectively. (L) Schematic diagram of CCR2 antagonist (INCB3344)-mediated inactivation of CCL2-CCR2 signaling to promote TMZ sensitivity of GBM cells. Experiments of C-K were independently used at least three different GAFs. Data in B-D, I, F, H, and J-K were mean±SEM. p values from the Pearson correlation test (B), one-way ANOVA (D and F,), and two-tailed student-t-test (H and J-K). * *P* < 0.05, ** *P* < 0.005, *** *P* < 0.001, **** *P* < 0.0001, ns= no significant.

### CCL2-CCR2 signaling activates the ERK1/2 expression in GBM cells to potentiate TMZ resistance

The CCL2-CCR2 axis was reported to reduce the response of TMZ treatment of glioma cells by activating intracellular G-protein receptors mediated downstream AKT signaling ^30^. We first investigated the differentially expressed genes of U87 and A172 cells after different conditioned cultures for three days. Gene set enrichment analysis revealed that the upregulated genes were significantly enriched in the positive regulation of the MAPK pathway (M11151:GO_POSITIVE_REGULATION_OF_MAPK_CASCADE, NES = 1.614 for U87 and 1.369 for A172, adjusted *P* < 0.05) (Figure 5A), which were known to be over-activated in treatment-resistant malignancies ^31,32^. Furthermore, WB analysis identified the upregulated expression of the phospho-ERK1/2 in the context of acquired anti-apoptosis of GBM cells following GAF-conditioned culture, whereas the expression JNK and p38 MAPK remained the same (Figures 5B and 5C). Patient-derived GBO models also expressed increased phospho-ERK1/2 protein after coculturing with GAFs (Figures S5A and S5B). Consistent with this interpretation, blocking the expression of MEK of GBM cells by trametinib, a potent, safe, and orally available MEK inhibitor^33^, during GAF coculture significantly increased the death rates of GBM cells with acquired chemoresistance. Meanwhile, trametinib alone could not ameliorate the acquired chemoresistance of GBM cells (Figures 5D and 5E). We next studied whether inhibiting MEK of GBM cells by trametinib in the conditioned culture system would also attenuate GAF-mediated chemoresistance. Our results showed that trametinib could decrease the TMZ resistance of GBM cells (Figures 5F and 5G; Figures S5D and S5E). Blocking the expression of ERK of GBM cells by another oral inhibitor ERK1/2, ravoxertinib (GDC-0994) ^34^, which resulted in phosphorylated-ERK1/2 downregulation, also led to dramatic cell death of resistant GBM cells (Figures 5H and 5I).

**Figure 5.**
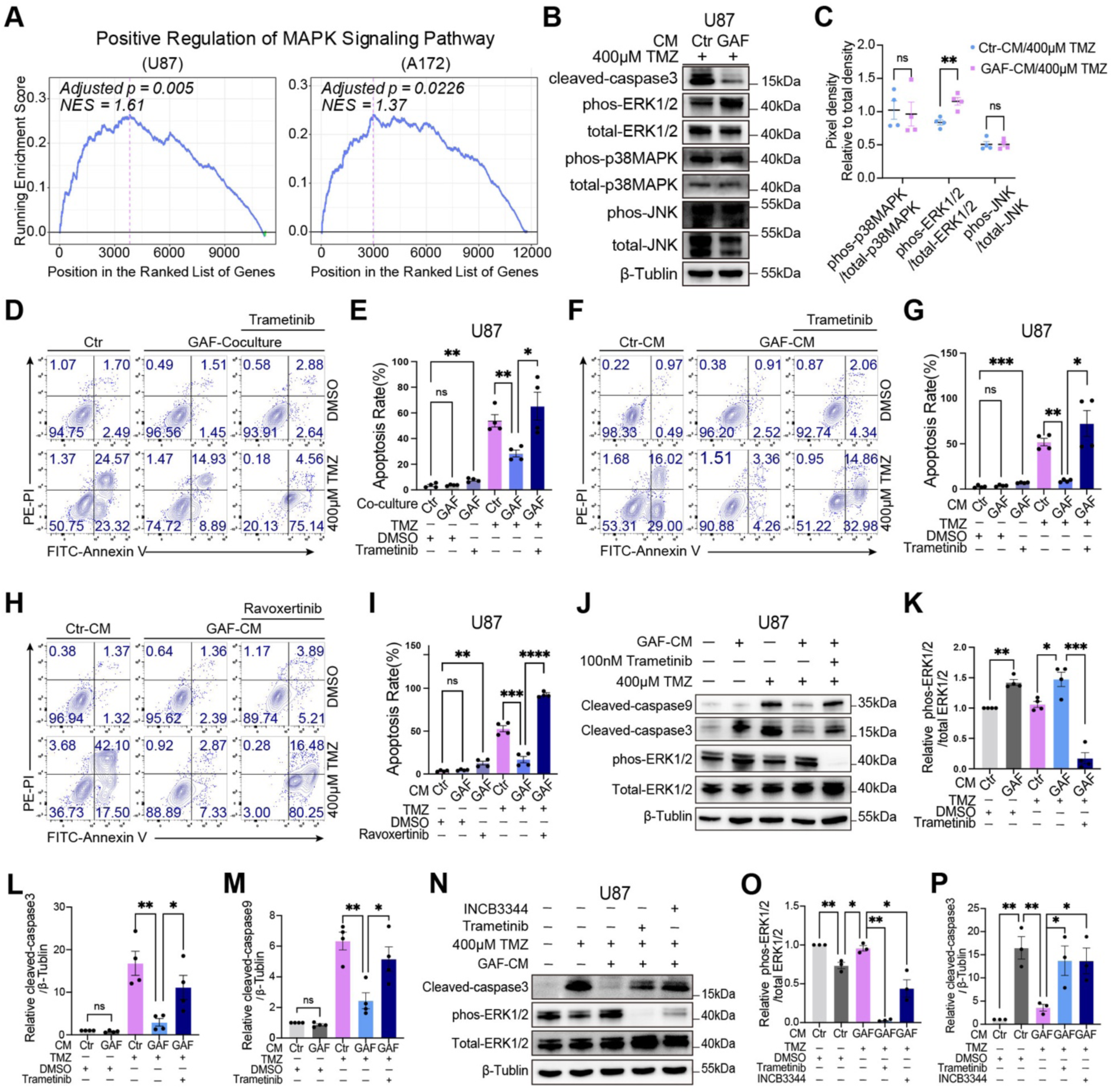
CCL2-CCR2 signaling activates the ERK1/2 expression in GBM cells to potentiate TMZ resistance. (A) Gene Set Enrichment analysis of differentially expressed genes on the MAPK pathway in glioma cells (U87 and A172) cultured in control medium or GAF conditioned medium. Adjusted *p* value, the estimated probability that a set with a given Normalized Enrichment Score represents a false positive finding (n = 6 per group). (B) WB analysis of three key molecules, such as phosphorylated ERK1/2(Thr202/Tyr204) and total ERK1/2, phosphorylated p38MAPK and total p38MAPK, and phosphorylated JNK1/2/3 and total JNK1/2/3, of MAPK signaling pathway of U87 cells cultured with Ctr-CM or GAF-CM for 48 hours following TMZ treatment respectively (C) Quantification of protein levels (4 different GAFs were used for this experiment). (D) Representative flow cytometry plots of apoptotic analyses of U87 cells following TMZ treatment for 48 hours. U87 cells were pretreated with Trametinib (100nmol/L) or DMSO for 6 hours, followed by U87 cells or GAFs coculture for 24 hours. (E) Quantification of apoptotic GBM cells (n = 4 per group). (F) Representative flow cytometry plots of apoptotic analyses of U87 cells following TMZ treatment for 48 hours. U87 cells were pretreated with Trametinib (100nmol/L) or DMSO for six hours, then cultured with Ctr-CM or GAF-CM for 48 hours. (G) Quantification of apoptotic GBM cells (n = 4 per group). (H) Representative flow cytometry plots of apoptotic analyses of U87 cells. U87 cells were pretreated with Ravoxertinib (20µmol/L) or DMSO for six hours, then cultured with Ctr-CM or GAF-CM for 48 hours. (I) Quantification of apoptotic GBM cells (n = 4 per group). (J) WB analysis of cleaved-caspase9, cleaved-caspase3, and phosphorylated ERK1/2 (Thr202/Tyr204) and total ERK1/2 in U87 cells with indicated treatment. (K-M) Quantification of protein levels (4 different GAFs were used for this experiment). (N) WB analysis of cleaved-caspase3 and phosphorylated ERK1/2 (Thr202/Tyr204), and total ERK1/2 in U87 cells with indicated treatment. (O-P) Quantification of protein levels (3 different GAFs were used for this experiment). Data in C, E, G, I, K-M, and N-P were mean±SEM. p values from empirical phenotype-based permutation tests and adjusted using the False Discovery Rate (FDR) procedure (A), two-tailed student-t-test (C, E, G, I, K-M, and N-P). * *P* < 0.05, ** *P* < 0.005, *** *P* < 0.001, **** *P* < 0.0001, ns = no significant.

Next, WB analysis suggested that the cleaved-caspase 3 and 9 in GBM cells cultured with GAF-conditioned media decreased dramatically under TMZ treatment, while MEK inhibition induced upregulation of those two apoptosis-related proteins in TMZ-resistant GBM cells (Figures 5J-L). Due to the activated ERK1/2 of resistant GBM cells, MEK inhibition revealed a dramatic rescue effect with downregulated expression of phospho-ERK1/2 (Figure 5M). To further investigate whether GAF-induced MEK1/2-ERK1/2 axis activation was mainly associated with GAF-derived CCL2, we conducted WB analysis to detect the level of phospho-ERK1/2 of GBM cells with INCB3344 treatment. Notably, our data indicated that in line with the rescue effect of trametinib, INCB3344 could significantly decrease the expression of phospho-ERK1/2 of GBM cells with acquired chemoresistance to improve their TMZ sensitivity, suggesting that GAF-derived CCL2 promotes TMZ resistance of GBM cells by activating the ERK1/2 expression (Figures 5N-P).

### Pharmacological inhibition of CCR2 or MEK1/2 enhances TMZ efficacy in patient-derived GBO models

Tumor organoids represented a useful platform for recapitulating the key features of parent tumors, which are valuable ex vivo models for testing new therapeutic and patient-specific treatment strategies in preclinical studies ^35^. We conducted ex vivo experiments to investigate whether GAFs promoted chemoresistance of patient-derived GBOs through CCL2-mediated ERK1/2 activation. Firstly, we successfully established patient-derived GBO models according to a previously reported method (Figure 6A), which was potent for rapidly generating GBOs ^36^. Our sphere-like GBOs grew in a suspended manner in the medium (Figure S6A), which maintained similar tissue architectures and cellular morphologies compared to their parent tumors (Figures 6B and 6C; Figures S6B and S6C). Immunofluorescence staining displayed the existence of tumor cells (GAFP+, Oligo2+, and Sox2+) and non-tumor cells in GBOs, such as macrophages (IBA1+) and GAFs (FN1+), which reserved the cellular heterogeneity of the parent tumor (Figures 6D and S6D). Thus, it demonstrated that our GBOs were credible tools imitating human GBMs for ex vivo experiments. Next, we cocultured GAFs and GBOs to test whether GAFs could promote the TMZ resistance of GBOs by detecting the expression of the cleaved-caspase 3. We found that GBOs following GAF stimulation exhibited less cleaved-caspase 3 signaling under TMZ treatment compared to the GBOs without GAF coculture (Figure 6E), which was consistent with our in vitro and in vivo findings. Importantly, we discovered a significantly increased expression of the cleaved-caspase 3 in TMZ-resistant GBOs in the condition of plus INCB3344 or trametinib targeted therapies (Figures 6E and 6F). To further elucidate whether ERK1/2 activation was the critical molecular mechanism rendering TMZ-resistant GBOs, IHC analysis revealed that GAFs increased the phospho-ERK1/2 expression in GBOs. Contrarily, INCB3344 or trametinib-treated TMZ-resistant GBOs showed significantly decreased phospho-ERK1/2 (Figure 6G). These results demonstrated that GAFs promoted the chemoresistance of GBM through CCL2-mediated ERK1/2 activation.

**Figure 6.**
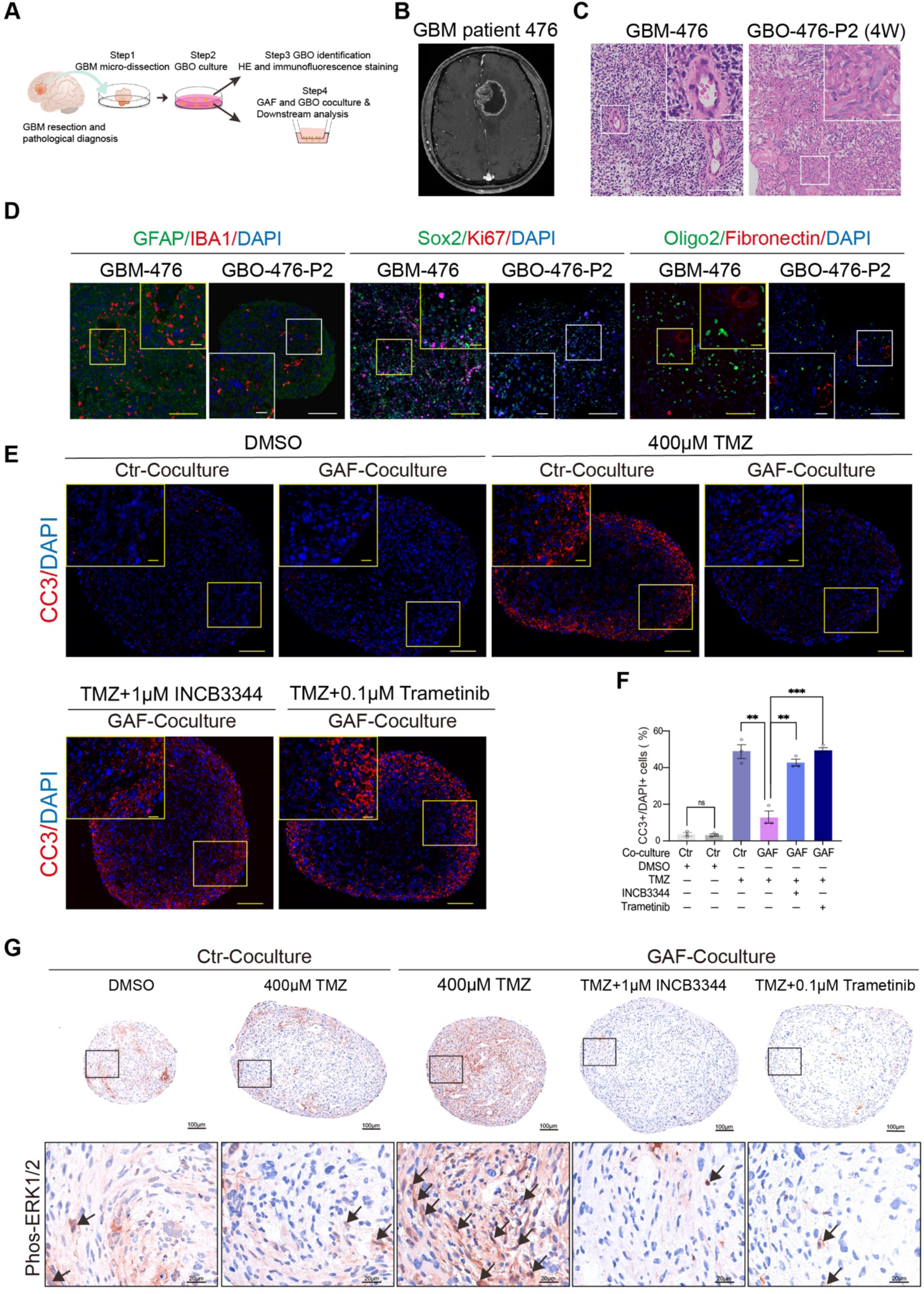
Pharmacological inhibition of CCR2 or ERK1/2 enhances TMZ efficacy in patient-derived GBO models. (A) Schematic diagram of GBO establishment and following experiments. (B) T1-weighted contrast-enhanced MRI images of patient 476. (C) Representative hematoxylin-eosin (HE) staining images of GBM tissue and corresponding GBO-476 (Passage 2). (D) Immunofluorescence images of representative markers of GBM in parent GBM-476 and GBO-476, such as GFAP, Oligo2, Sox2, IBA1, FN1, and Ki67, showing that GBO-476 maintained cellular heterogeneity and proliferation. Scale bars for the original images and the magnified images were 100µm and 20µm respectively. (E) Representative immunofluorescence images of cleaved-caspase3 (CC3) in GBOs following indicated treatment. Scale bars for the original images and the magnified images were 100µm and 20µm respectively. (F) Quantification and comparison of the fraction of CC3+/DAPI+ cells in each GBO group (n = 3 different patient-derived GBOs per group). (G) Representative IHC images of phos-ERK1/2 in GBOs following indicated treatment. Scale bars for the original images and the magnified images of (**C-E**, and **G**) were 100µm and 20µm respectively. Data in (F) were mean±SEM. p values from two-tailed student-t-test (F). ** P < 0.005, *** P < 0.001, ns = no significant.

## Discussion

In our study, we isolated GAFs from gliomas using our optimized cell culture method and demonstrated that GAF-secreted CCL2 stimulated CCR2 on GBM cells, which promoted activation of the ERK1/2 expression to potentiate chemoresistance, thus ameliorating TMZ-induced tumor cell apoptosis. INCB3344 or trametinib administration contributed to reversing GBM cells from a TMZ-resistant to a TMZ-sensitive phenotype via blocking the CCL2-CCR2 axis or MEK1/2-ERK1/2 pathway, respectively.

Although many solid tumors harbor CAFs in their microenvironment ^11,20^, CAFs residing in glioma have been identified recently by single-cell analyses ^15,16^. Based on this crucial finding, we further provided experimental evidence demonstrating that GAFs were present in both IDH-mutant human gliomas and IDH-wildtype GBMs and held a function of mediating chemoresistance. Due to a lack of consensus-specific CAF markers, most previous studies isolated CAFs either by utilizing growth competition advantage and differential adherence of CAFs for serial trypsinization in the cell culture approach, or differential gravitational gradient between CAFs and epithelial cells using centrifugation ^8,9^. However, in previous reports and our prior experience, the cell type remains highly heterogeneous using the traditional cell culture method even after multiple passages, thus hindering the effective purification ^37^. This may be explained by the low fraction of CAFs in gliomas, and the epithelial-to-mesenchymal transition (EMT) phenomenon in some tumor cell populations. Using neural stem cell medium, as shown in our results, significantly improves the separation of CAF-like cells from tumor cells within glioma tissue, and in all our cases allows isolation of purified CAFs at Passage 0, which enables us to check the authenticity of CAFs in gliomas and study their biological function with sufficient passages before senescence.

The origin of GAFs should be elucidated in future investigations. While CAFs were identified as cancer stromal cells exhibiting elongated spindle-shaped morphology and markers of activated fibroblasts including fibronectin, α-SMA, Vimentin, and CD90, heterogeneity of CAFs has been frequently reported ^6^. In our RNA analysis, we found pronounced expression of the Asporin (ASPN) gene in GAF samples. A similar result was observed in CAFs derived from prostate cancer ^38^, but ASPN was expressed at a significantly lower level in breast cancer and gastric cancer CAFs that we included. Cancer stem cells undergoing profound epithelial-to-mesenchymal transition were reported to be the potential origin of CAFs ^39^. As IDH is a decisive gene in classifying the diagnosis and prognosis of diffuse gliomas ^3^, future studies detecting whether GAFs harbor this tumor-specific mutation would help judge the cell origins of GAFs. Moreover, CAFs from the same cancer type could also exhibit differences. One such example was that a subset of CAFs in pancreatic ductal adenocarcinoma showed diminished α-SMA expression, and instead secreted IL6 and other inflammatory mediators ^20^. In addition, single-cell analysis of GBM reveals the presence of distinct molecular profiles between earlier-stage GBM CAFs and late-stage GBM CAFs ^15^. The proximity between GAFs, CMASCs, and MSCs in their transcriptomic profile suggests that GAFs and CMASCs may share a common ancestor with MSCs, or possibly arise from MSCs, as MSCs could develop into CAFs within the tumor microenvironment ^23^. However, recent evidence suggested that resident fibroblasts in the CNS could proliferate and participate in the pathological response to inflammation ^13^. Therefore, it is also reasonable to hypothesize that GAF could arise from resident fibroblasts in the CNS.

CAFs have been shown to contribute to tumor aggressiveness by supporting tumor growth, facilitating invasion, recruiting tumor-protective niche, and tampering with anti-tumor immune response ^39^. However, the role of GAFs in the treatment resistance of tumors has not been described. The resistance of malignant gliomas to first-line chemotherapy agent TMZ was believed to be one of the major factors behind the poor patient outcomes of this disease. No significant survival benefit was observed in patients receiving TMZ beyond 6 cycles, suggesting that gain of chemoresistance during treatment ^40^. Most previous reports attributed the chemoresistance of gliomas to the heterogeneity and plasticity of the tumor cells ^41,42^. Here, we expand the mechanism of the TMZ resistance of GBMs by showing that GAFs, an exogenous factor, participate in this process via secreting chemokine CCL2. Our data align with another finding that pericyte promotes GBM chemoresistance through CCL5-CCR5 signaling-mediated enhanced DNA damage repair ^43^. Taken together, non-tumor cells in the glioma microenvironment play complicated roles in controlling the fate of tumors. Future studies should consider those non-tumor cells if any therapeutic strategy is about to be developed. Given that IDH-mutant gliomas also contain fewer GAFs, we should elucidate why, compared to GBMs, their quantities decrease significantly. CAFs were known to communicate with cancer cells via growth factors, cytokines, or chemokines ^5^. The CCL2 could reduce the response of TMZ treatment of glioma cells through intracellular G-protein receptors mediated downstream of AKT signaling activation and promoting glycolysis ^30^. Despite the CCL2 could activate G-protein receptors to upregulate multiple intracellular oncogenic pathways, such as JAK/STAT3, PI3K/AKT, and MAPK pathways, previous reports found that CCL2-induced chemoresistance of various tumors was associated with PI3K/AKT signaling, few studies mentioned the contribution of MEK1/2-ERK1/2 pathway activated by CCL2 ^26^. Specific molecular mechanisms regulating CCL2-mediated ERK1/2 activation should be discussed in the future.

It reported that continuous activation of the MEK1/2-ERK1/2 pathway was associated with acquired chemoresistance of GBM and gastric cancer ^44,45^. In non-small cell lung cancers, CAFs could counteract the effect of tyrosine kinase inhibitors on EGFR+ tumors by secreting HGF, which activated MET Proto-oncogene and downstream signaling, including PI3K-AKT and MAPK ^8^. In our study, we successfully inhibited the protective role of GAFs in the GBM cells by blocking the interaction of GAF-GBM cells via the CCR2 antagonist INCB3344. In addition, inhibiting MEK by trametinib potently decreased TMZ resistance of GBM cells resulting from GAF stimulation. Thus, we demonstrated that CCL2 secreted by GAFs coincided with upregulation of the ERK signaling pathway which could abrogate the intrinsic apoptosis in the glioma cells induced by TMZ treatment. Our study found a distinct mechanism that GAF was an extraneous factor promoting continuous ERK1/2 activation in GBM cells. Moreover, CCL2 secreted by GAFs induced chemoresistance of GBM cells through MEK1/2-ERK1/2 signaling activation, rather than previously reported CCL2-mediated PI3K/AKT activation ^26^. We also successfully validated our findings of the relationship between GAFs and chemoresistance of GBMs by ex vivo GBO models, which encouraged us to take advantage of the patient-derived organoids to assess the chemoresistance-promoting role of GAFs in GBMs and conduct preclinical investigations to target GAF-GBM interplay. Our study suggested that counteracting the GAF-GBM interplay may be a novel druggable candidate to improve the efficacy of chemotherapy for GBM patients.

## Limitations of the study

We acknowledge that using our hospital scRNA-seq data of glioma samples would strengthen the evidence of GAFs growing in low-grade gliomas and GBMs. Although we found cytokine CCL2 secreted by GAFs could promote TMZ resistance, further testing of specific mechanisms mediating abundant CCL2 secretion is warranted. Finally, our data strongly indicate the cytokine CCL2 induced ERK1/2 activation in mediating the TMZ resistance, however, future extensive in vitro and in vivo experiments are necessary to detect the efficacy of GAF-targeted therapy in restraining GBM recurrence.

## Methods

### GAF establishment

We discovered an optimized approach in isolating the glioma-associated fibroblasts from fresh biopsies based on traditional method of culturing primary fibroblasts^17^, which was likely to produce pure GAFs in early cell generations during fibroblasts establishment. Processing began within 1 hour after samples collection. Biopsies were mechanically minced with sterile scissors into fragments of less than 1 mm^3^ and enzymatically digested with 20 mg/mL papain (Worthington #20132), 0.7mg/mL hyaluronidase (Sigma #V900833) and 0.7mg/mL collagenase-I (Sigma #C0130) for 20 minutes in a water bath of 37℃, mixing tissue suspension gently with a pipette every 5 minutes. After digestion, red blood cells were ruptured by the red blood cell lysis buffer (Beyotime #C3702). To eliminate cancer stem cells, we adopted culture medium screening method, primary miscellaneous cells were first resuspended in neural stem cell medium consisting of DMEM/F12(Gibco #A4192001) supplemented with 2% B27(Gibco #17504044), b-FGF (20ng/mL, Peprotech #100-18B), EGF (20ng/mL, Peprotech #AF-100-15), streptomycin-penicillin (100U/mL, Gibco #15140122) for 24 hours at 37℃. Then, the supernatant was sucked out and the dish was gently tapped and washed with PBS to remove non-adherent cells and tissue debris. subsequently, cells were cultured in MesenCultTM Proliferation Kit (Human) (STEMCELL #05411) to purify the primary GAFs. In most cases (95%), the pure fibroblast-like cells were successfully isolated in passage 0. The pure GAFs population grew to a confluence of 90% usually costing approximately 15 days in R10 plastic plates. All experiments used GAFs within passage 10 to avoid cell senescence and genetic alterations.

### GBO generation and culturing

According to the previously developed methodology to generate GBO by F. Jacob et al ^36^, we successfully established GBOs to conduct the ex vivo experiments. Briefly, resect fresh en bloc GBM tissue without heavy cauterization and suction, then transiently store them in a pre-cooling Hibernate A medium (Thermo Fisher Scientific #A1247501). Tissue processing started within 2 hours after tissue removal. Carefully cut GBM tissue into 0.5-1mm small pieces in a sterilized glass dish using sharp-edged microsurgical scissors. Incubate tumor pieces with the red blood cell lysis buffer (Beyotime #C3702) for 10 minutes at room temperature to lyse red blood cells. Wash tumor pieces three times with DMEM/F12 (Thermo Fisher Scientific #11320033), and then transfer the dissected tumor pieces to non-adherent 6-well culture plates with 4 ml of GBO medium per well. Components of GBO medium: 235ml of DMEM/F12, 235ml of neurobasal medium (Thermo Fisher Scientific #21103049), 5ml of MEM-NEAAs solution (Thermo Fisher Scientific #11140050), 5ml of GlutaMAX supplement (Thermo Absin Fisher Scientific #35050061), 5 ml of penicillin-streptomycin, 5 ml of N2 supplement (Thermo Fisher Scientific #17502048), 10 ml of B27 minus vitamin A supplement (Absin #abs9272), and 125μl of human recombinant insulin (Absin #abs42225219). Culture GBO on an orbital shaker rotating at 100 rpm. GBO should become round and cell-dense morphology in about 2 weeks. Take GBOs with a diameter of 0.5-1mm at different time points to validate the cellular consistency between GBOs and parental GBMs, and to conduct chemotherapeutic experiments. GBO should be cut into smaller pieces under the microscope once its volume reaches 1-2 mm to alleviate necrotic cell death within the core. All experiments used GBOs within passage 2 to avoid the tumor microenvironment alteration, for example, cell types.

### Cell-conditioned culture, cell coculture, and cell apoptosis assay

Conditioned-culture and coculture assays were conducted to investigate the effect of GAFs on the chemoresistance of GBM cells.

Cell-conditioned culture: a). Preparing for conditioned medium (CM): 1×10^5^ GAFs or 1×10^5^ GBM cells (U87 or U251) were seeded in the 6-well plastic plate respectively and were cultured in DMEM 10%FBS for 24 hours. Subsequently, supernatants underwent centrifugation to discard cell debris at 1000x g for 5 minutes and were collected as a CM for the following conditioned culture. b). Conditioned culture: 1×10^4^ glioma cells were evenly seeded on slides in 12-well plastic plates. Add 1mL of CM into the plastic well and renew the CM every 12 hours for indicated days.

Cell coculture: Experiments were performed within the SPLInsert 6 well transwell apparatus (SPL life sciences #37006) which contains 0.1µm polyethylene terephalate (PET) membrane of the upper chamber. GAFs (2×10^4^) and GBM cells (U87 or U251) (2×10^4^) were respectively cultured in the upper or lower chamber for the indicated time. d). Cell apoptosis assay: To investigate the protective effect of GAFs on glioma cells, DMSO or TMZ (400µmol/L for U87, and 800µmol/L for U251) (Selleckchem #S1237) will be administrated to induce apoptosis of glioma cells for 48 hours. Apoptosis analysis was performed using the Annexin V-FITC/PI kit (4Abiotech #FXP018) for flow cytometry on a Beckman cytoflex. Hochest33258 staining kit was used to analyze apoptosis of glioma cells and images were photographed by Olympus BX63 Microscope. Image J was used for cell counting.

### GBO-GAF coculture and treatment

Use GBOs with similar sizes under microscopy to conduct each assay. Coculturing GAFs and GBOs in the SPLInsert 6 well transwell apparatus with the medium of DMEM/F12 containing 1% FBS. For the control group, the same patient-derived GBOs were cultured in the upper and lower chambers respectively for three days. For the experimental group, GAFs (2×10^4^) and GBOs were cultured in the upper and lower chambers respectively for three days. Then, organoids were subjected to 0.1% DMSO, or 400 µM TMZ, or a combination of TMZ plus 100 nM trametinib or 1µM INCB3344 for five days. Add 2 ml of 4% (vol/vol) formaldehyde solution to fix GBOs for half an hour and conduct immunofluorescence staining of Cleaved-caspase3 and IHC analysis of phos-ERK1/2.

### Primary glioma specimens

Fresh glioma samples were collected into sterile cold streptomycin-penicillin PBS buffer within 10 minutes after resection. All human tissues utilized in the present study were approved by the Ethics Committee on Biomedical Research, West China Hospital of Sichuan University (No.2021.1319), and attained written informed consent from patients or their legal guardians. Histopathological diagnosis of gliomas was performed by two neuropathologists in the department of pathology according to the fifth edition of the World Health Organization (WHO) Classification of CNS Tumors. Cryopreserved and formalin-fixed, paraffin-embedded glioma samples were stored in Liquid nitrogen or at 4℃. All procedures were performed by the principles of the Helsinki Declaration.

### GAF identification through scRNA-seq of gliomas

The scRNA-seq data of primary GBMs and LGGs was acquired from the GSE182109 GEO series published by Abdelfattah N et al^18^. We removed low-quality single cells expressing < 500 genes or containing > 20% mitochondrial or > 50% ribosomal transcripts, as well as potential single-cell doublets identified by DoubletFinder (with default settings), based on the filtered Cell Ranger feature counts aligned to GRCh38 (human), as described by Abdelfattah N et al. We also removed genes expressed in fewer than 3 single cells. Only samples were included with over 1000 remaining cells in downstream analysis. We modified the clustering steps due to recent updates in Seurat. Robust Principal Component Analysis (RPCA, provided by Seurat) of the top 2000 most variable genes was first used to integrate multiple samples into one Seurat dataset on 30 RPCA components. Further batch correction and dimensional reduction were conducted using Harmony. Next, Seurat was used to find cell clusters using the Leiden algorithm on the Harmony reduction variables. Seurat FindAllMarkers (min. pct = 0.10, logfc. threshold = 0.10) were also used to identify marker genes for each cluster. Then, we used the SingleR to annotate the identity of each single cell based on BlueprintEncodeData, HumanPrimaryCellAtlasData reference datasets and a reference set consisting of RNA-sequencing data acquired from Sequence Read Archive (SRA), which included previously reported CNS-metastasis associated stromal cells (PRJNA510710), CAFs (GSE106503, GSE112350, GSE195832), MSCs (GSE105145, GSE73610), skin fibroblasts (GSE113957), pericytes (GSE104141), endothelial cells (GSE159851), astrocytes (GSE179882), and microglial cells (GSE133432). Finally, we conducted CNV analysis using infercnv with macrophages and T cells as normal reference. Only 10000 cells were randomly sampled with stratification by clustering were shown in Fig. S2D due to plot rendering capacity of our device.

### Glioma cells culture

GBM cell lines U87 and U251 were brought from the China Center for Type Culture Collection (CCTCC, Wuhan, China). A172 was bought from Procell (#CL-0012). The lineages of GBM cells were identified by Short Tandem Repeat (STR) analysis. Tumor cells were cultured in Dulbecco’s modified Eagle’s medium with high Glucose (DMEM, High glucose, BIOEXPLORER # B1101-001) supplemented with 10% fetal bovine serum (FBS, ExCell Bio #FSP500) in 5% CO2 in a humidified incubator at 37°C.

### GAFs RNA sequencing and pre-processing

Total RNA was extracted from tissue and cells using Trizol reagent (Invitrogen, USA). mRNA was purified from total RNA using poly-T oligo-attached magnetic beads. Sequencing libraries were generated using NEBNext® UltraTM RNA Library Prep Kit for Illumina® (NEB, USA) following the manufacturer’s recommendations. Each sequencing library was sequenced on Illumina Hiseq 4000 to generate 6 gigabases of 150bp paired-end reads. Clean reads were mapped to the GRCh37 human reference genome and counted using STAR (2.6.0c).

### GAFs cell identity analysis

We inferred the cell type identity of the patient-derived cells (PDCs) using CellO. Briefly, the algorithm quantified log transcript per million (TPM) of genes and predicted the probability of the sample being each cell type using isotonic regression (IR) or cascaded logistic regression (CLR) model. For this analysis, we used the pre-trained models provided by the developers, which were based on the comprehensive data set comprising transcriptomic data from diverse cell types. We also predicted the cell type identity of control glioma cell lines and the aforementioned reference transcriptome set. To focus on the most specific results, we only kept cell type labels where the probability of any sample exceeds 0.5 and that of any sample fell below 0.3. To understand if the patient-derived cells were malignant, we used CNVkit to estimate genomic copy number variations (CNV) with RNA-seq data. We used the copy number and RNA-seq count matrix of the TCGA-LGG GBM cohort to generate CNV-expression correlation coefficients for this analysis. The tumor-adjacent brain and the skin fibroblasts were supplied as normal controls. Log2 ratio under –0.7 or above 0.7 and weight greater than 100 were used as a cutoff to call a CNV event.

### Functional analysis of differentially expressed genes after conditioned culture

DESeq2 was used to call differentially expressed genes between control cells and conditioned-cultured cells. ClusterProfiler package ^46^ was used to conduct gene set over-representation (ORA) and enrichment analysis (GSEA) of the differentially expressed genes with adjusted p-value under 0.05.

### Bioinformatic analyses of human gliomas from the TCGA database

The transcriptomic profile data of TCGA-GBM and TCGA-LGG samples and associated clinical information were downloaded using TCGA-biolinks ^47^. R package EPIC (Estimate the Proportion of Immune and Cancer) was used to estimate CAF fraction from the transcriptomic profile data of TCGA glioma samples^48^. Samples with a CAF fraction over 0.5 were dropped in further analysis because these samples were likely to be fibrotic scar tissue (n = 5).

### Immunofluorescence and immunohistochemistry (IHC) staining

Immunofluorescence and IHC staining were performed according to standard procedures. Cells were seeded on slides with a confluence of 50% in 24-well plates. slides were washed twice using phosphate-buffered saline (PBS) and then permeabilized with PBS containing 0.25% Triton X-100 for 20 minutes. Blocking was conducted with PBS plus 10% normal goat serum for 90 minutes, followed by incubation with the primary antibody in PBS with 3% normal goat serum overnight at 4°C. Primary antibody included α-SMA (Proteintech #55135-1-AP, 1:300), Fibronectin (BD Biosciences #610077 1:500), FAP (Abcam #ab53066, 1:200), Oligo2 (Millipore #ab9610, 1:300). Slides were incubated with fluorophore-conjugated secondary antibodies at a dilution of 1:1000 for 90 minutes at room temperature. Washing slides with PBS 3 times and finally mounting slides with Prolong TM anti-fade reagent with DAPI (Invitrogen #P36935). Paraffin-embedded mice brain sections, GBM tissues, and GBO tissues were cut into 3µm thickness and stored at 4°C. Tissue sections were incubated with the primary antibody: Cleaved-caspase3 (Cell Signalling Technology #9661, 1:200) and phos-ERK1/2 (Cell Signalling Technology #4370S, 1:400) overnight at 4°C, and stained with the SABC kit (Boster #AR1020) and DAB Chromogenic Substrate Kit (Boster #AR1022) according to the standard protocols. For tissue immunofluorescence staining, tissue sections were incubated with the primary antibody: CD105 (Abcam #ab169545 1:400), fibronectin (BD Biosciences #610077 1:200), overnight at 4°C. Quantification of DAB positive area was performed using ImageJ with the plugin of IHC profiler. Stained slides were photographed by Olympus BX63 Microscope. All slides stained with the same antibodies were digitalized using uniform acquisition settings.

### qRT-PCR

Total RNA was extracted by the cell total RNA isolation Kit (FOREGENE #RE-03113) according to the operating manual. cDNA synthesis was performed with PrimeScriptTM RT reagent Kit (Takara #RR037B), and quantitative assays were performed in triplicate using SYBR Premix Ex TaqTM II (Takara #RR820A) in Bio-Rad iQ5 system. Relative gene expression was normalized to β-actin. The primer sequences were listed in Table S2.

### Western blotting and antibodies

Protein electrophoresis was performed according to standard procedures. Primary antibodies were prepared at a 1:1000 dilution and were then probed with HRP-linked secondary antibody (1:10000). Information of primary antibodies was as follows: Fibronectin (BD Biosciences #610077), α-SMA (Proteintech #55135-1-AP), Cleaved-caspase3 (Cell Signaling Technology #9661), Cleaved-caspase9 (Cell Signaling Technology #9509), Bax (Millipore #ABC11), β-Tublin (Millipore #MAB3408), β-Actin (Cell Signaling Technology #3700), GAPDH (Proteintech #CL594-60004), Phospho-ERK1/2(Thr202/Tyr204) (Cell Signaling Technology #4370S), total-ERK1/2 (Cell Signaling Technology #4696), Phospho-p38 MAPK (T180/Y182) (Abclonal #AP1165), total-p38 MAPK (Abcam #ab308333), Phospho-JNK1-T183/Y185+JNK2-T183/Y185+JNK3-T221/Y223 (Abclonal #AP0276), total-JNK1/2/3 (Abclonal #A4867), CCL2/MCP1 (Abclonal # A23288).

### ELISA assay

The supernatants were prepared as follows: 1×10^5^ GAFs or U87, U251 cells were cultured in 6-well plastic plates respectively with DMEM 1%FBS for 72 hours then supernatants from different groups were collected. The supernatants were stored in aliquots at –80°C. CCL2 concentration in supernatants was normalized by cell number and analyzed by ELISA (R&D) according to the manufacturer’s instructions.

### Flow cytometry

For fibroblast-specific biomarkers expression analysis, 5×10^4^ alive GAFs or glioma cells were incubated with PE Mouse anti-Human CD105 (2µg/mL, BD Biosciences #560839) and APC Mouse anti-Human CD90 (2µg/mL, BD Biosciences #559869) for 30 minutes at room temperature. Wash cells with pre-cooling PBS twice. Flow cytometry was performed with a Beckman cytoflex. All files of flow cytometry were processed with FlowJo v10.10.0 software.

### Orthotopic GBM xenografts and treatment

All animal experiments were approved by the Institutional Animal Care and Use Committee (IACUC) of West China Hospital, Sichuan University complying with the Guide for the Care and Use of Laboratory Animals. BLAB/c female Nude mice with a mean age of 6 weeks were purchased from Beijing Vital River Laboratory Animal Technology (China). Briefly, for control groups, 5×10^4^ U87 cells expressing luciferase were transplanted into the corpus callosum of the right frontal lobe of five Nude mice aged 6 to 8 weeks. Likewise, 5×10^4^ U87 cells plus 1×10^4^ GAFs were implanted into the same location for the experiment group of five Nude mice. To determine whether GAFs were able to confer chemoresistance to GBM cells, following bioluminescence detection of tumors at day 20, tumor-bearing mice were treated with TMZ (40mg/kg, i.p) at day 20 to 24. Post-therapy bioluminescence was conducted at the day 25 and day 32. Tumor volume was monitored through bioluminescence using IVIS Spectrum apparatus (Caliper Life Sciences) after i.p injection of D-luciferin 150mg/Kg (GOLDBIO #LUCK-1G). Mice were sacrificed when manifested more than 20% of body weight loss or neurological symptoms, such as hemiparesis or somnolence.

## QUANTIFICATION AND STATISTICAL ANALYSIS

For ORA the p values were calculated using hypergeometric distributions. These p values were adjusted in the Benjamini–Hochberg approach. Prism 9.0 software was employed in Statistical analysis. All grouped data were presented as box plots in figures. All quantitative data were means ± SEM. Unpaired Student-t test or one-way analysis of variance (ANOVA) analysis was performed for experiments with two groups or more than two groups, respectively, except for the comparison between the estimated CAF fractions of different glioma groups where we used the Wilcoxon signed-rank test. For comparison between the estimated CAF fractions of primary and recurrent tumors, we used the paired Wilcoxon signed-rank test. Kaplan-Meier survival curves were generated using Prism software, and the log-rank test was performed to assess statistical significance between groups. A two-sided *p* value lower than 0.05 was considered significant.

## Data and code availability

Raw sequencing data is available at Genome Sequence Archive for Humans (https://ngdc.cncb.ac.cn/gsa-human/s/OnfgtAhQ) with accession HRA002403 and could be accessed under regulation of the GSA-Human Data Access Agreement. The code used for analyses is available at https://github.com/sxz-ivan/GAF-analysis. All data are available in the main text or the supplementary materials.

## Author contributions

M.R.Z., M.N.C., and Y.H.L. investigated and designed the study. M.R.Z., S.X.Z., S.L.C., Y.B.Y., Y.Z.H., W.H.L performed all experiments. M.R.Zuo., S.X.Z., Y.B.Y., S.L.C., W.C.Y., T.F.L., Z.H.W., W.T. F., and Y.F.X. collected and analyzed all data. N.C. examined the diagnosis of human gliomas. M.R.Z., S.X.Z., M.N.C., Y.H.Z., Y.Y., Q.M., and Y.H.L. supervised the study. All authors wrote, reviewed, edited, and approved the publication of the original paper.

## SUPPLEMENTAL INFORMATION

### Supplementary Figures

**Figure S1.**
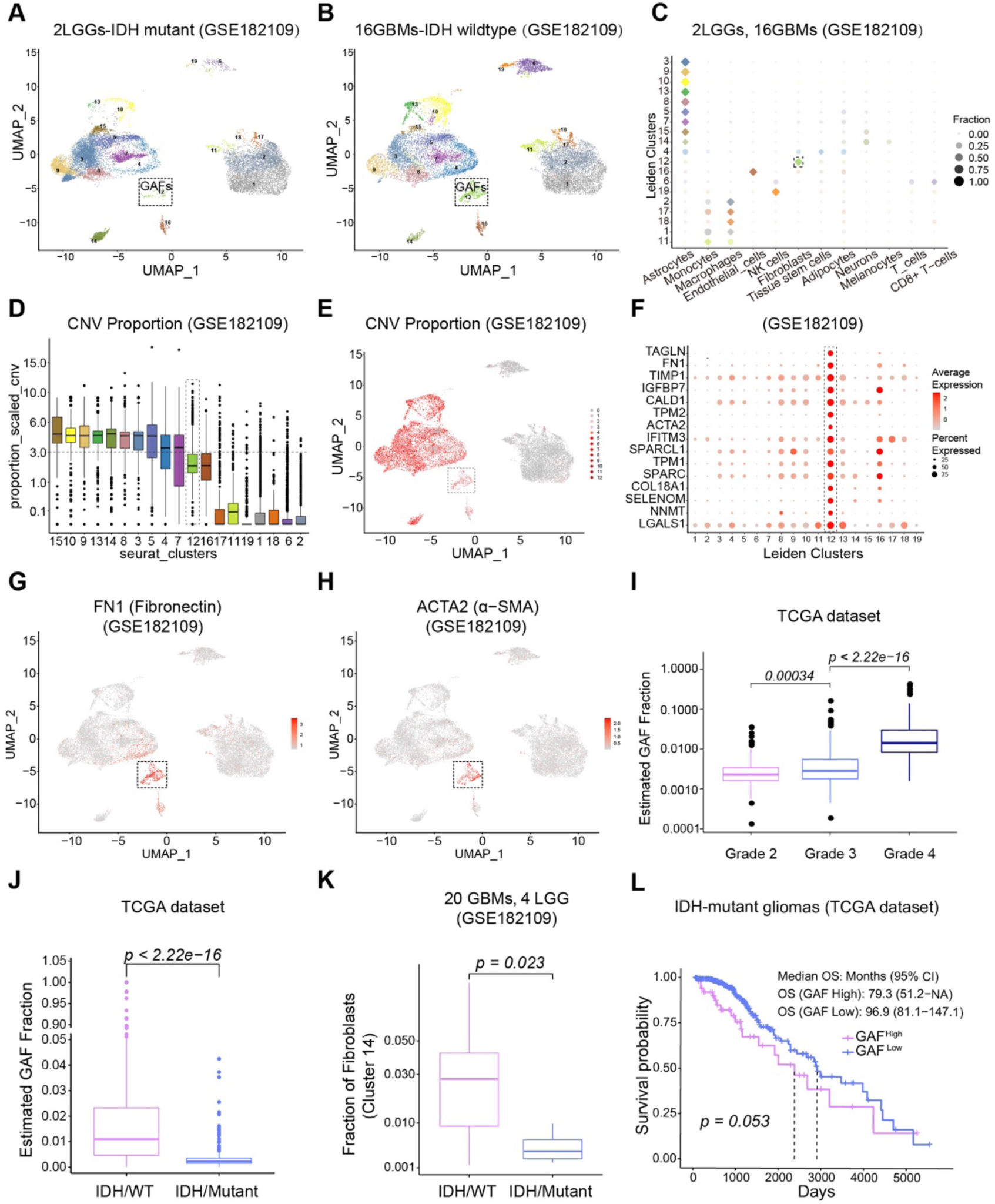
scRNA-seq reveals that GAFs exist in distinct grades of gliomas and are associated with clinical characteristics of gliomas, related to Figure 1. (A and B) UMAP plot presentation of cells clustered by Seurat and their SingleR annotation based on the GSE182109 scRNA-seq dataset. (A) 2 LGGs and (B) 11 primary GBMs and 5 recurrent GBMs. (C) SingleR annotations of cells in each cluster (squares represent annotations with the highest fraction in each cluster. (D) CNV analysis of each cell type in scRNA-seq using infer CNV. (E) UMAP plot presentation of CNV of each cell. (F) Bubble plot showing the average expression of cluster genes for each cluster. The bubble size indicates the percentage of cells expressing a marker, and the color indicates the expression level. (G and H) Feature expression plots of Fibronectin (FN1) and ACTA2 (α-SMA) expression of each cell cluster. (I) The relationship of estimated GAF fraction and WHO grade of gliomas from the TCGA database. (J) The estimated fraction of GAFs in IDH-wildtype (IDH/wt, n=218) and IDH-mutant (IDH/mut, n=398) gliomas from the TCGA cohort computed by the EPIC R package. (K) The fraction of GAFs in GBMs (IDH/wt, n=20) and lower-grade gliomas (IDH/mut, n=4) in the GSE182109 scRNA-seq dataset. (L) Kaplan–Meier survival analysis of overall survival of patients with IDH-mutant gliomas stratified by their respective GAF level from the TCGA database. Data in I-L were mean±SEM. p values from the two-tailed Wilcoxon ranked sum test (I-K) and log-rank test (L).

**Figure S2.**
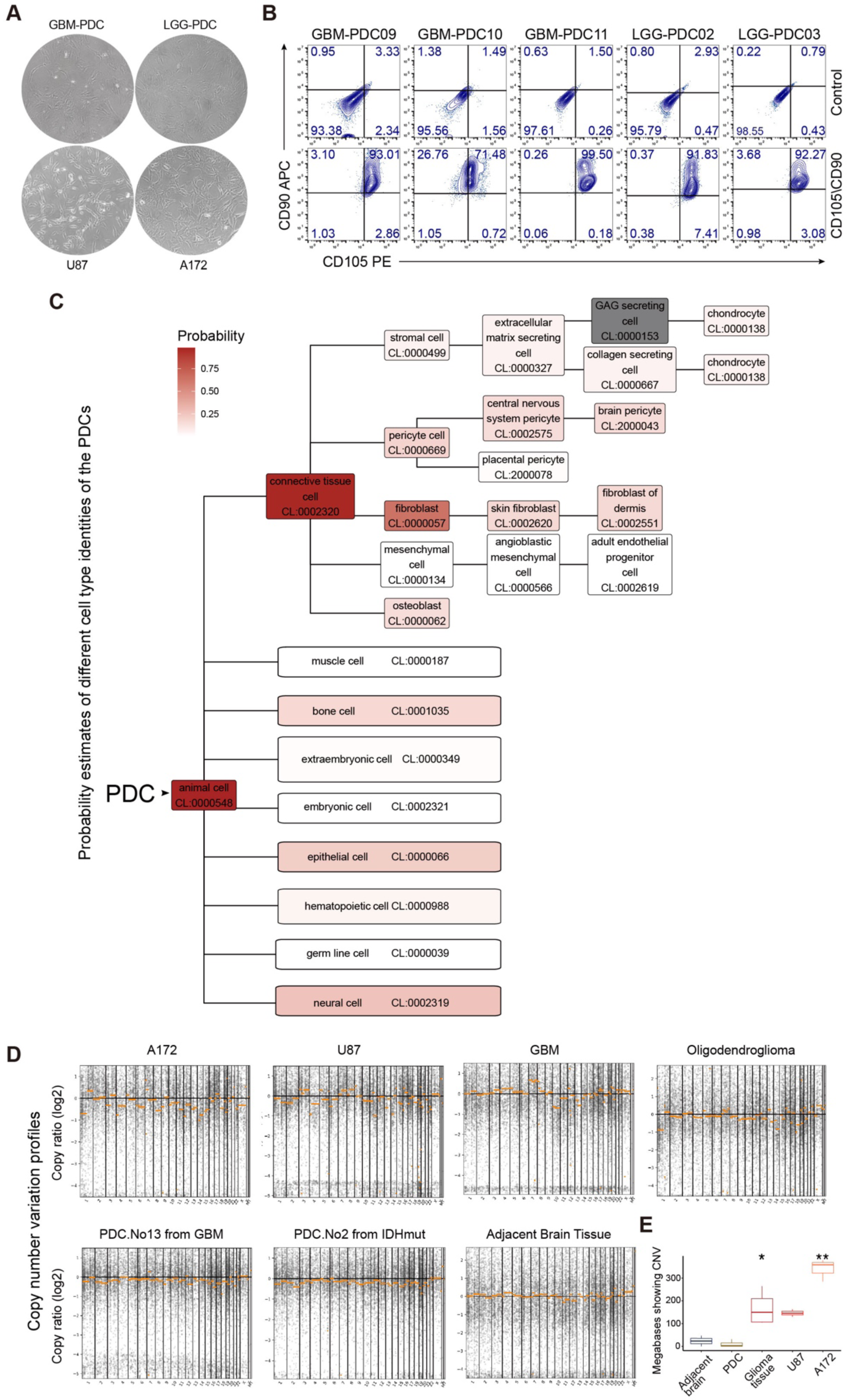
Primary culture and identification of patient-derived GAFs from human gliomas, related to Figure 2. (A) Representative morphological images of PDCs isolated from GBM (GBM-PDC) and lower-grade glioma (LGG-PDC) under a light microscope. The morphology of U87 and A172 is the negative control. Magnification, 200x. (B) Proportions of PDCs (PDC02,03,09,10,11) expressing CAF markers (CD105, CD90) analyzed by flow cytometry (FCM). The upper panel was negative control without primary antibodies staining. The bottom panel was stained with antibodies. (C) Probability estimates of different cell type identities of the PDCs in the Cell Ontology vocabulary system with the pre-trained isotonic regression (IR) model. (D) Representative images of estimated CNV profiles of each cell and tissue sample. (E) Comparison of estimated CNV profiles of each cell and tissue sample. Data in (E) were mean±SEM. p values from the two-tailed Wilcoxon-ranked sum test (E). * *P* < 0.05, ** *P* < 0.005.

**Figure S3.**
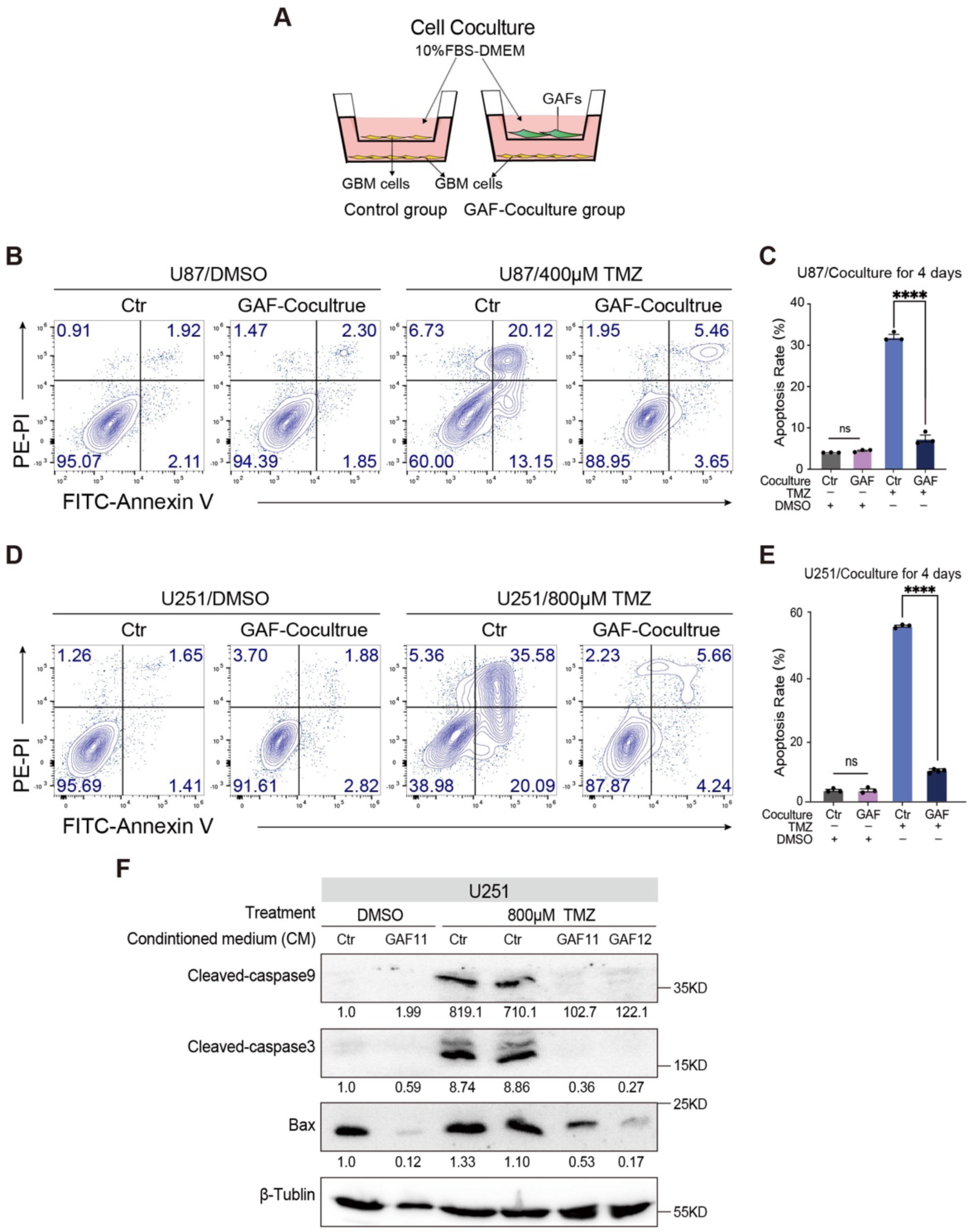
GAFs protect GBM cells from TMZ-induced apoptosis, related to Figure 3. (A) Schematic diagram of 6-well transwell coculture of GAFs and GBM cells. DMSO or TMZ was added into the lower chamber to treat GBM cells after coculture for indicated hours. Apoptosis analyses were performed 48 hours after the treatment. (B) Representative images of the percentage of apoptosis of U87 cells cocultured with GAFs or GBM cells for 96 hours following TMZ treatment. (C) Quantifications of apoptotic GBM cells (n = 3 per group). (D) Representative images of the percentage of apoptosis of U251 cells cocultured with GAFs or GBM cells for 96 hours following TMZ treatment. (E) Quantifications of apoptotic GBM cells (n = 3 per group). (F) WB analysis of cleaved-caspase9, cleaved-caspase3, and Bax in U251 cells following indicated treatment. Experiments in B-E were independently used at least three different GAFs. Data in C and E were mean±SEM. p values from two-tailed student-t test (C and E). **** p < 0.0001, ns = no significant.

**Figure S4.**
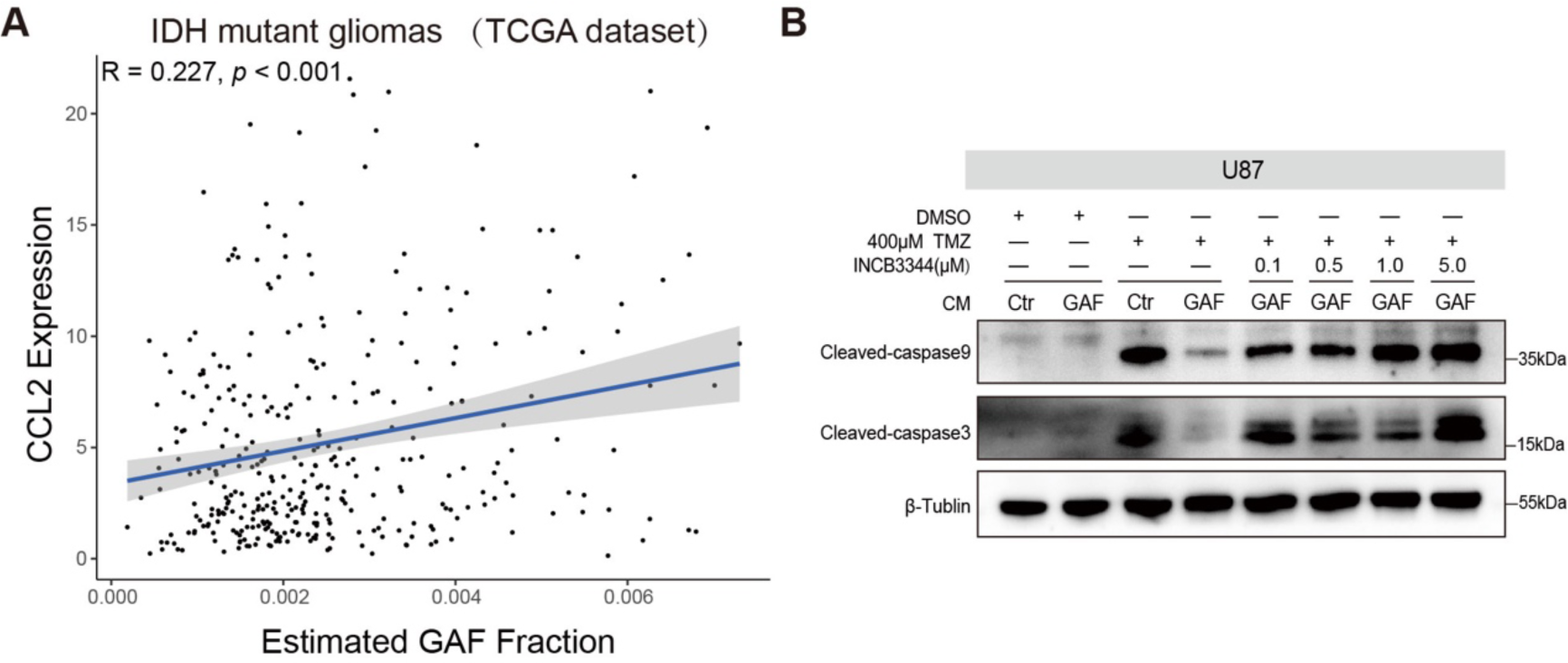
GAF promotes TMZ resistance of GBM cells via cytokine CCL2, related to Figure 4. (A) Correlation analysis of CCL2 expression and estimated GAF fraction of lower-grade gliomas from the TCGA database (n=398). (B) To find out an appropriate concentration of INCB3344 to antagonize CCR2, GBM cells were cultured with Ctr-CM or GAF-CM supplemented with a concentration gradient of INCB3344 (0.1µmol/L, 0.5µmol/L, 1µmol/L, 5µmol/L). Then, WB analysis of cleaved-caspase9, and cleaved-caspase3 in GBM cells was conducted following indicated treatment. Data in A were mean±SEM. p values from the Pearson correlation test (A).

**Figure S5.**
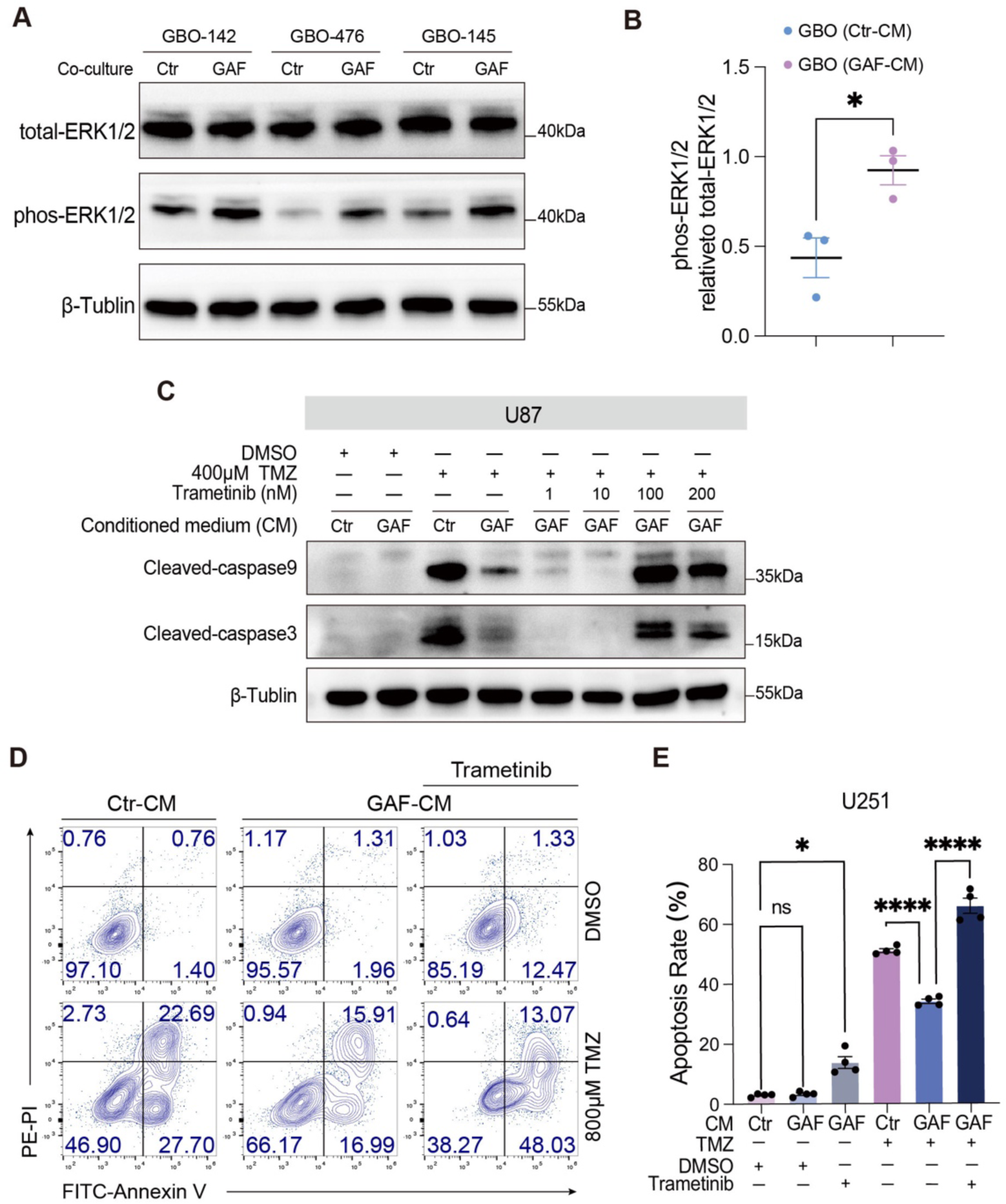
CCL2-CCR2 signaling activates the ERK1/2 pathway in GBM cells to potentiate TMZ resistance, related to Figure 5. (A) Expression of phosphorylated ERK1/2 and total ERK1/2 in patient-derived GBO models following coculture with GAFs (B) Quantification of the protein levels (n = 3 different patient-derived GBOs per group). (C) To find out an appropriate concentration of Trametinib to inhibit ERK1/2, GBM cells were. cultured with Ctr-CM or GAF-CM supplemented with concentration gradients of Trametinib (1nmol/L, 10nmol/L, 100nmol/L, 200nmol/L). Then, WB analysis detecting cleaved-caspase9 and cleaved-caspase3 in GBM cells was conducted following the indicated treatment. (D) Representative flow cytometry plots of apoptotic analyses of U251 cells following TMZ. treatment for 48 hours. U251 cells were pretreated with Trametinib (100nmol/L) or DMSO for six hours, then cultured with Ctr-CM or GAF-CM for 48 hours. (E) Quantification of apoptotic GBM cells (n = 4 per group). Data in (B and E) were mean±SEM. p values from two-tailed student-t-test (B and E). * p < 0.05, **** p < 0.0001, ns = no significant.

**Figure S6.**
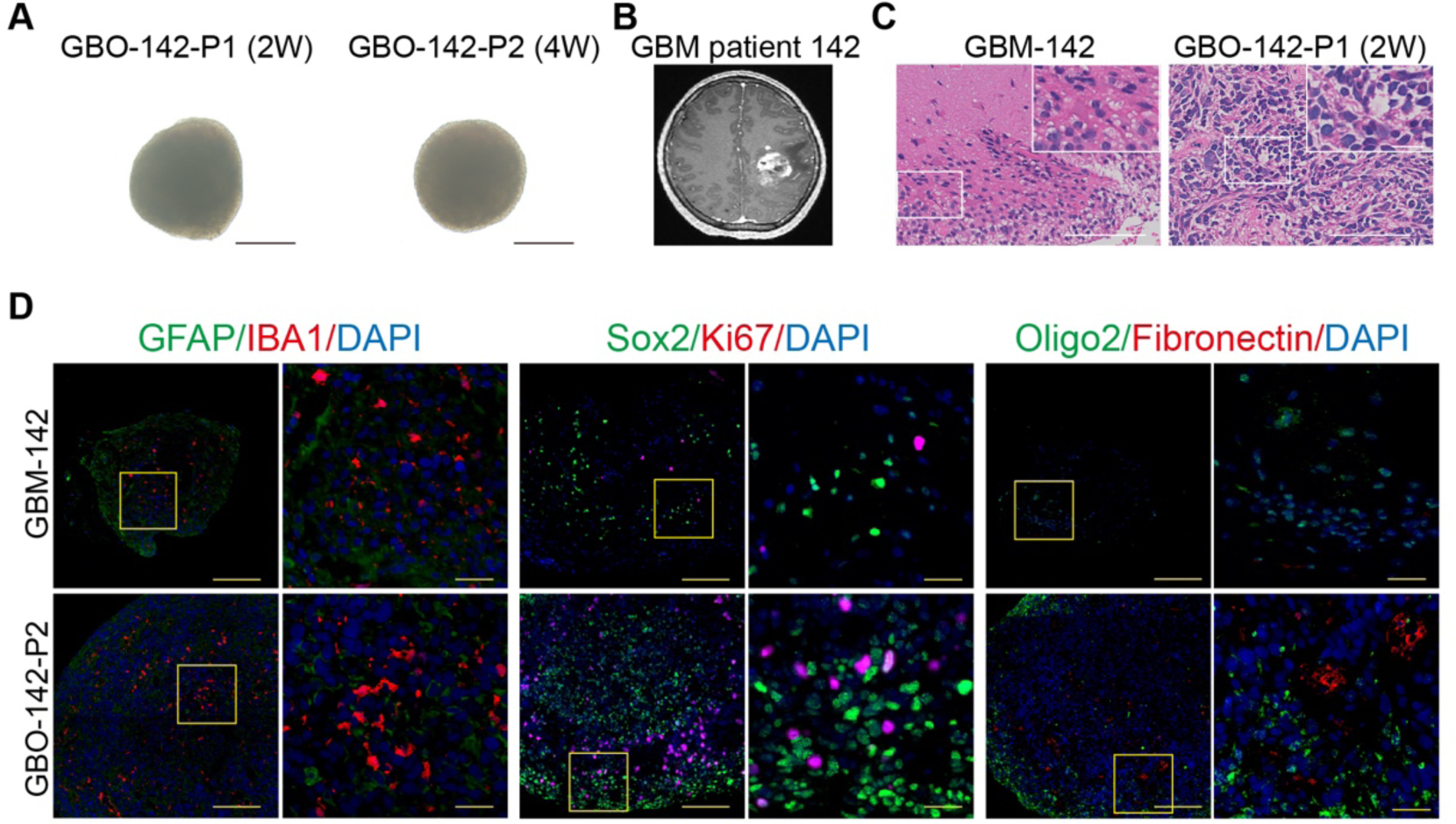
Patient-derived GBOs establishment, related to Figure 6. (A) Representative images of GBO-142 (Passages 1 and 2) under a microscope. Scale bar, 500µm. (B) T1-weighted contrast-enhanced MRI images of patient 142. (C) Representative HE images of GBM tissue and corresponding GBO-142 (Passage 2). (D) Immunofluorescence images of representative cellular markers of GBM in parent GBM-142 and GBO-142. Scale bars for the original images and the magnified images of (C and D) were 100µm and 20µm respectively.

**Supplementary Table 1.** The patient demographic information of matching GAFs and GBOs.

| Number | Pathology | Location | Experiments |
| --- | --- | --- | --- |
| GAF01 (PDC01) | Anaplastic oligodendroglioma, grade III | Frontal lobe | RNA-seq, FCM, qPCR, WB, IF, Apoptosis analysis |
| GAF02 (PDC02) | Anaplastic oligodendroglioma, grade III | Frontal lobe | RNA-seq, FCM, qPCR, WB, IF, Apoptosis analysis |
| GAF03 (PDC03) | Oligodendroglioma, grade II | Frontal lobe | RNA-seq, FCM, qPCR, WB, |
| GAF04 (PDC04) | Oligodendroglioma, grade II | Frontal lobe | RNA-seq, qPCR, WB, |
| GAF05 (PDC05) | Oligodendroglioma, grade II | Frontal lobe | RNA-seq, qPCR, WB, |
| GAF06 (PDC06) | Recurrent anaplastic oligodendroglioma, grade III | Frontal lobe | RNA-seq, WB, |
| GAF07 (PDC07) | Glioblastoma, grade IV | Frontal lobe | RNA-seq |
| GAF08 (PDC08) | Glioblastoma, H3K27M mutant | Thalamus | RNA-seq, FCM, qPCR, WB, Apoptosis analysis, Animal study |
| GAF09 (PDC09) | Recurrent glioblastoma, grade IV | Frontal lobe | RNA-seq, FCM, qPCR, WB, Apoptosis analysis |
| GAF10 (PDC10) | Glioblastoma, grade IV | Parietal lobe | RNA-seq, FCM, qPCR, WB, IF, Apoptosis analysis, Animal study |
| GAF11 (PDC11) | Recurrent diffuse astrocytoma, grade II | Frontal lobe | RNA-seq, FCM, WB |
| GAF12 (PDC12) | Glioblastoma, grade IV | Temporal lobe | WB, Apoptosis analysis, IF |
| GAF13 (PDC13) | Glioblastoma, grade IV | Frontal lobe | WB |
| Glioma01 | Recurrent diffuse astrocytoma, grade II | Frontal lobe | RNA-seq |
| Peritumoral brain01 | / | Frontal lobe | RNA-seq |
| Glioma02 | Glioblastoma, grade IV | Frontal lobe | RNA-seq |
| Peritumoral brain02 | / | Frontal lobe | RNA-seq |
| Glioma03 | Anaplastic oligodendroglioma, grade III | Frontal lobe | RNA-seq |
| Glioma04 | Recurrent glioblastoma, grade IV | Temporoparietal lobe | RNA-seq |
| GBO-142 | Primary glioblastoma, grade IV | Temporal lobe | WB, HE, IF, IHC, Apoptosis analysis |
| GBO-476 | Primary glioblastoma, grade IV | Frontal lobe | WB, HE, IF, IHC, Apoptosis analysis |
| GBO-145 | Primary glioblastoma, grade IV | Parietal lobe | WB, HE, IF, IHC, Apoptosis analysis |

**Supplementary Table 2.**
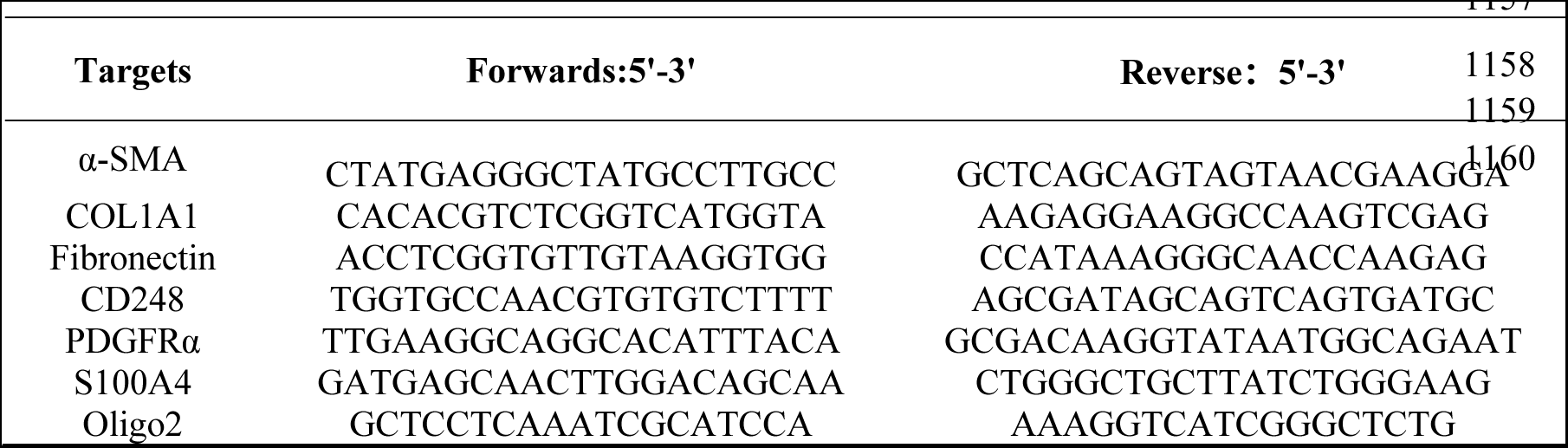
The primer sequences for qRT-PCR.

